# A prenatal window for enhancing spatial resolution of cortical barrel maps

**DOI:** 10.1101/2024.03.15.585193

**Authors:** Mar Aníbal-Martínez, Lorenzo Puche-Aroca, Gabriele Pumo, M. Pilar Madrigal, Luis M. Rodríguez-Malmierca, Francisco J. Martini, Filippo M. Rijli, Guillermina López-Bendito

## Abstract

Precise mapping of peripheral inputs onto cortical areas is required for appropriate sensory processing. In the mouse primary somatosensory cortex, mystacial whiskers are represented in large barrels, while upper lip whiskers are in smaller, less defined barrels. Barrel size and definition of these two functionally distinct barrel maps are believed to be determined by the type of whisker input and peripheral receptor density. However, spontaneous activity and transcriptional programs at prenatal developmental stages can influence somatosensory map development independently of sensory experience. Thus, the mechanisms defining distinct barrel field territories, including their size and definition, remain poorly understood. Here, we show that prenatal ablation of mystacial whiskers remap somatosensory cortical territories resulting in enhancement of the functional and anatomical definition of upper lip whisker barrels. These changes do not result from alterations in receptor type but rather stem from thalamic upper lip input-receiving neurons adopting a mystacial-like transcriptional profile. Our results unveil a regulated prenatal mechanism within the thalamus that maps available somatosensory input to ensure sufficient cortical barrel size and functional spatial resolution for sensory processing, irrespective of peripheral receptor type and density.

## INTRODUCTION

Sensory systems are represented in the primary sensory areas of the brain, structured into both anatomical and functional maps^1–4^. The spatial precision of sensory maps varies among species according to the ethological relevance of the modality or the intra-modal sensory branch^5^. This precision is achieved through the accuracy of point-to-point innervation along the ascending sensory pathway. In rodents, which heavily rely on the somatosensory modality, a prominent cortical area is devoted to processing stimuli from facial whiskers. Within the mouse primary somatosensory cortex (S1), information from facial whiskers is processed in two juxtaposed subfields: the postero-medial barrel subfield (PMBSF) and the antero-lateral barrel subfield (ALBSF). The PMBSF receives input from the mystacial whiskers which provide diverse and specialized information to S1^6,7^. The ALBSF receives input from the upper lip whiskers which are considered to play a secondary role in sensory processing^8^. While each barrel corresponds to a single whisker in both regions, the degree of clustering of incoming thalamocortical axons varies, leading to notable anatomical distinctions between the two territories. While PMBSF barrels are characterized by their large size and sharp borders, ALBSF barrels are smaller and have poorly defined borders^9^. These differences in intra-modal map organization parallel the distinct morphological characteristics of mystacial and upper lip whiskers: mystacial whiskers are long with large follicles, whereas upper lip whiskers are comparatively short with small follicles, albeit more abundant. Despite their clear anatomical and functional differences, we still lack a complete understanding of the mechanism underlying the construction of these two distinct barrel maps and to what extent they rely on the type of sensory receptors (mystacial versus upper lip). Indeed, recent data demonstrated that sensory receptor-independent mechanisms can also influence cortical barrel size by altering the patterns of activity in subcortical stations. For instance, increasing thalamic waves in the developing somatosensory thalamus of early blind mice results in larger barrels in the PMBSF of S1, even before the onset of active whisking^10,11^. Thus, the size of a sensory barrel map may not only depend on the type and number of sensory receptors but may also be regulated by intrinsic programs during development. By embryonically ablating mystacial whiskers and follicles, and the subsequent PMBSF representation, we generated mice in which the ALBSF barrels become larger and better-defined, without altering the size and number of upper lip follicles. This reorganization of the barrel maps occurs within a restricted prenatal time window and is guided by transcriptional programs operating intra-modally in the thalamus. Interestingly, these reorganized ALBSF barrels resemble normal PMBSF barrels at the morphological, molecular and functional levels, showing an enhanced spatial resolution upon tactile stimulation.

## RESULTS

### The size of barrel subfield areas can be adjusted prenatally

The size of the areas devoted to processing distinct whisker -mystacial versus upper lip- input information is thought to be determined by the corresponding type and density of sensory receptors on the mouse face^12,13^. We asked whether prenatal removal of a selected facial whisker type might have an impact on the cortical barrel areas and spatial distribution of the spared whiskers. We implemented a strategy whereby the mystacial whiskers in the whisker pad were cauterized unilaterally at embryonic day (E) 14 (named as embWPC), in both wild type and TCA-GFP mice^14^, in which thalamocortical projections are labeled with green fluorescent protein (GFP). At this stage, trigeminal nerve axons have just begun to target the principal trigeminal sensory nucleus (PrV) in the brainstem (**Figure S1A**), but PrV neuron axons have not yet entered the ventral posteromedial (VPM) thalamic nucleus^15–19^.

Analysis of the snout at E18 confirmed the specific ablation of the mystacial whiskers and follicles (**Figure 1A**), with a strong reduction of whisker pad-innervating primary trigeminal sensory axons targeting the PrV (**Figure S1B**). At early postnatal stages, the total areas of PrV, thalamic ventral posterior nucleus (VPN) and S1, including other somatosensory body representations apart from the snout, did not exhibit significant differences in embWPC, as compared to control mice (**Figures S2A-S2C**). In contrast, the size of the areas corresponding to mystacial and upper lip whiskers significantly rescaled in embWPC mice at all levels of the pathway, suggesting intra- modal plasticity of connectivity. For example, in S1, we found a 54% decrease in the PMBSF territory, and a 34% expansion of the ALBSF territory, a phenomenon already observed at P4 (**Figures 1B** and **S2D**). Notably, we found no discernible differences in the total upper lip volume, nor in the volume and number of upper lip follicles between control and embWPC mice both at E18 and P8 (**Figure 1A**; **Videos S1** and **S2**). Dye depositions in the cortical PMBSF or ALBSF of P8 embWPC confirmed a similar intra- modal remapping of the corresponding whisker pad input-receiving VPM (wpVPM) or upper lip input-receiving VPM (ulVPM) areas in the thalamic VPM^20^ (**Figure 1C**). Moreover, retrogradely labeled thalamocortical neurons within each VPM sub-territory maintained their expected point-to-point distribution, matching cortical topography (**Figure S3**).

**Figure 1.**
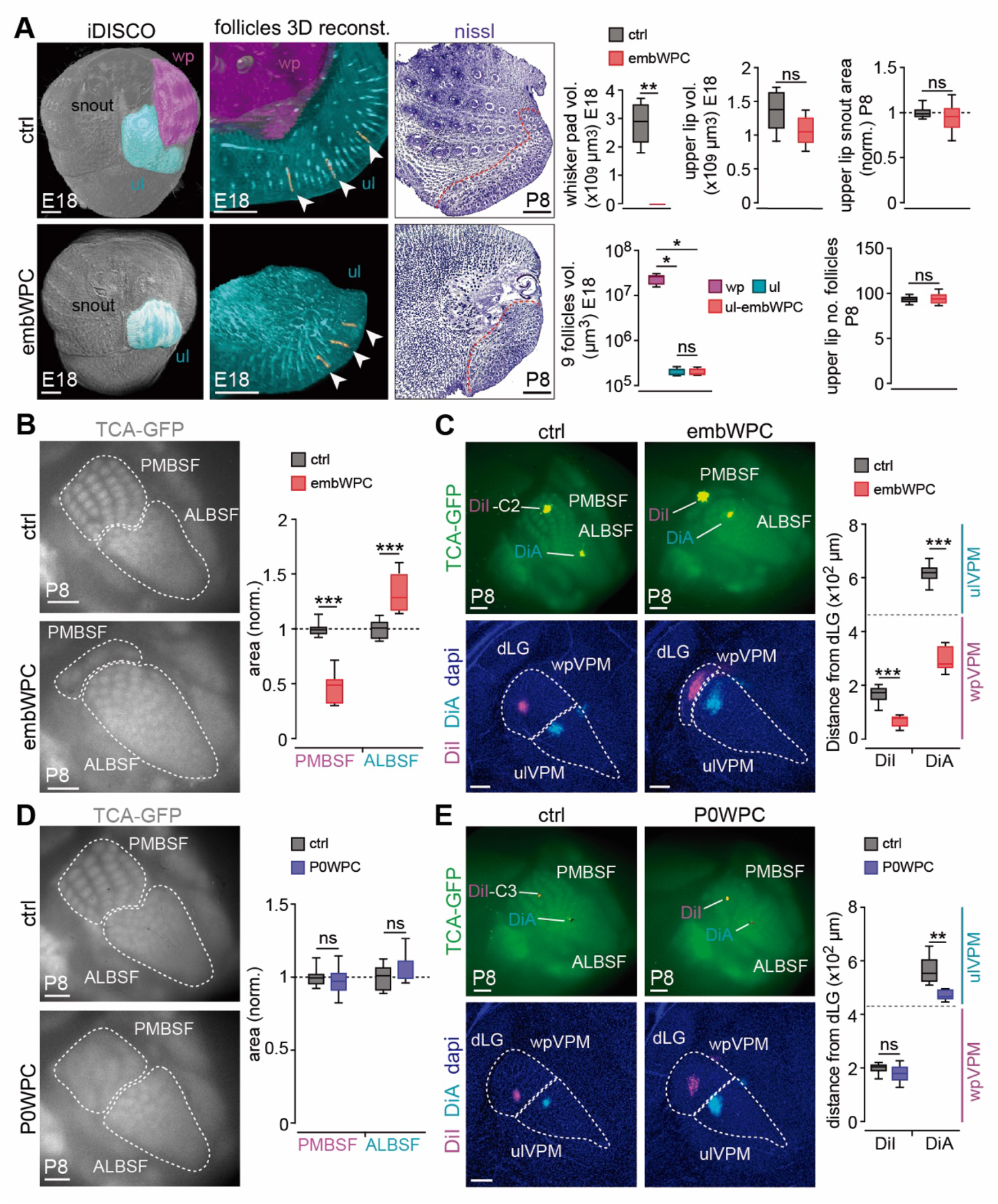
Whisker pad deprivation in embryos rescales barrel field areas and thalamocortical circuits. (A) Left and middle, iDISCO frontal snout views showing the whisker pad (magenta) and upper lip (cyan) areas, and lateral 3D-reconstructions of the follicles (yellow) of the snout at E18 in control and after whisker pad cauterization of the mystacial whiskers mice. Right, Nissl stainings of the snout at P8 in control and embWPC mice, red dashed line delimits upper lip area. Quantification of the data (iDISCO E18, n= 5 ctrl mice, n= 5 embWPC mice, whisker pad volume, unpaired two-tailed Student *t*-test; upper lip volume, unpaired two-tailed Student *t*-test. ns, p= 0.11; upper lip area, unpaired two-tailed Student *t*-test. ns, p= 0.32; upper lip 9 follicles volume, Mann-Whitney *U*-test. ns, p= 0.73. Nissl P8, n= 8 ctrl mice, n= 9 embWPC mice; upper lip number of follicles, unpaired two-tailed Student *t*-test. ns, p= 0.78). **(B)** Surface view of thalamocortical terminals (TCA-GFP+) in the cortical PMBSF and ALBSF in control and embWPC mice at P8. Quantification of the data (n= 8 ctrl mice, n= 9 embWPC mice, unpaired two-tailed Student *t*-test). **(C)** Upper panels, DiI and DiA crystal placements in the C2 cortical barrel and in the ALBSF, respectively, in control and embWPC at P8. Lower panels, backlabelled cells at the level of the wpVPM and ulVPM showing the relative displacement of the labelling in embWPC mice. Quantification of the position of backlabelled cells with respect to the distance to the dLG nucleus. The gray horizontal dashed line in the graph represents the separation between wpVPM and ulVPM (n= 6 ctrl mice, n= 6 embWPC mice, unpaired two-tailed Student *t*-test). **(D)** Surface view of thalamocortical terminals (TCA-GFP+) in the cortical PMBSF and ALBSF in control and P0WPC mice at P8. Quantification of the data (n= 8 ctrl mice, n= 9 P0PC mice, unpaired two-tailed Student *t*-test. ns, PMBSF, p= 0.55, ALBSF, p= 0.30). **(E)** Upper panels, DiI and DiA crystal placements in the C3 cortical barrel and in the ALBSF, respectively, in control and P0WPC at P8. Lower panels, backlabelled cells at the level of the wpVPM and ulVPM. Quantification of the position of backlabelled cells with respect to the distance to the dLG nucleus. The gray horizontal dashed line in the graph represents the separation between wpVPM and ulVPM (n= 5 ctrl mice, n= 5 P0WPC mice, unpaired two-tailed Student *t*-test. ns, p= 0.09). wp, whisker pad; ul, upper lip; E, embryonic; P, postnatal; embWPC, embryonic whisker pad cauterized; P0WPC, postnatal day 0 whisker pad cauterized; vol., volume; no., number; TCA-GFP, thalamocortical axons labelled with green fluorescent protein; PMBSF, postero-medial barrel subfield; ALBSF, antero-lateral barrel subfield; wpVPM, whisker pad recipient ventral posteromedial nucleus; ulVPM, upper lip recipient ventral posteromedial nucleus; norm., normalized. Scale bars, (A, left) 500 μm; (A, middle) 400 μm; (A, right) 1000 μm; (B), (C, top), (D) and (E, top) 500 μm; (C, bottom) and (E, bottom) 200 μm. Boxplots show the medians with the interquartile range (box) and range (whiskers). Bar graphs show the means ± SEM. ns, not significant. *p < 0.05, **p < 0.01, ***p < 0.001. See also Figures S1-S5; Videos S1 and S2.

Importantly, the changes of the cortical barrel field sub-territories depended on mechanisms only present before birth, since cauterization of the whisker pad at postnatal day 0 (P0), referred to as P0WPC, revealed no significant territorial differences in the PMBSF and ALBSF between control and P0WPC across the somatosensory stations (**Figures 1D** and **S4**). As expected, there were no re-arrangements of the thalamocortical axons in the P0WPC (**Figures 1E** and **S5**). Thus, there is a critical window in which the cortical ALBSF area can be significantly adjusted without altering upper lip receptors, suggesting that after this timepoint plasticity might arise from a different strategy to compensate for sensory loss.

### Functional rescaling of ALBSF in embWPC mice occurs before birth

Whisker pad stimulation *in vivo* at E18 can elicit activity in the developing PMBSF^21,22^, indicating that the periphery-to-cortex somatosensory pathway is already functional before birth. We asked whether the anatomical changes observed in embWPC at P8 were detectable at prenatal stages by assessing functional reorganizations. We carried out peripheral whisker pad and upper lip stimulations at E18 while recording cortical calcium activity in *Emx1^GCaMP6f^* transgenic mice, expressing GCaMP6f in cortical glutamatergic neurons^22^ and carrying unilateral whisker pad cauterization (**Figure 2A**).

**Figure 2.**
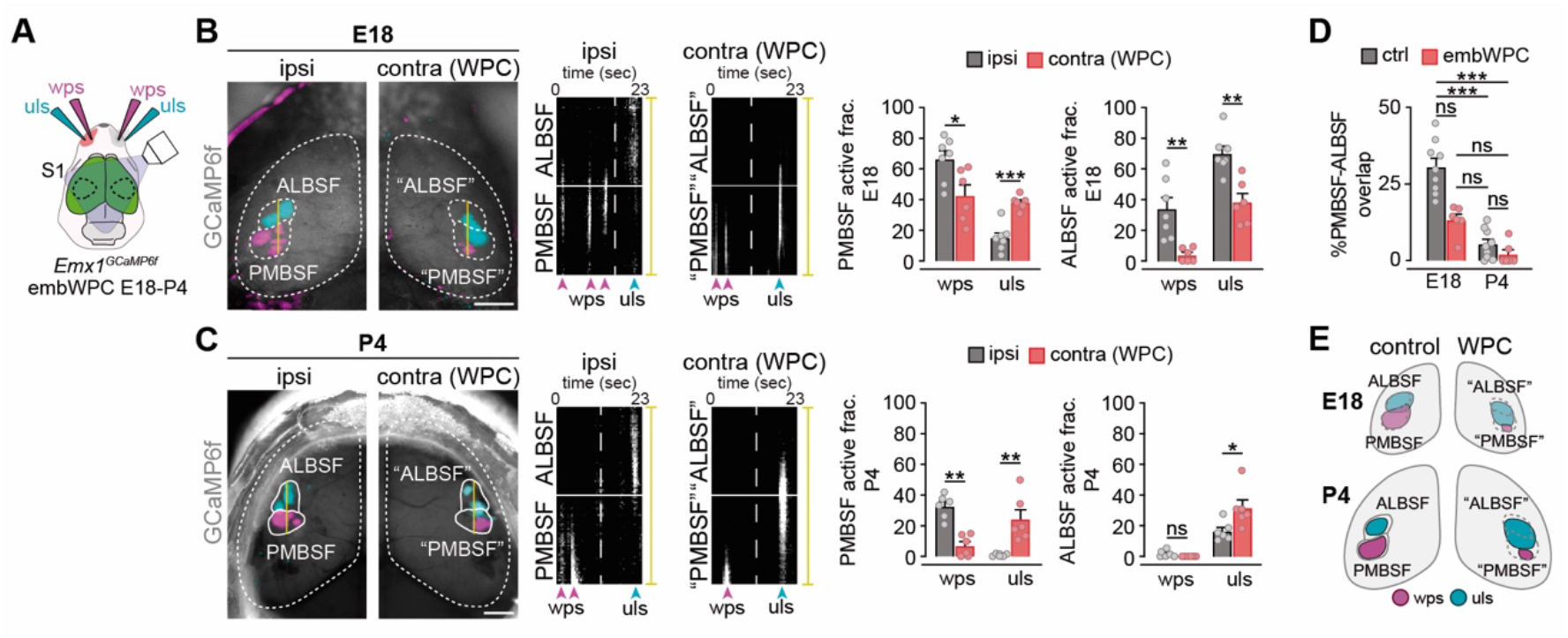
Functional rescaling of cortical barrel field territories in embWPC mice. **(A)** Schema representing the experimental paradigm. **(B)** Left, cortical evoked responses (GCaMP6f) to whisker pad (magenta) and upper lip (cyan) stimulations in control (ipsilateral) and whisker pad cauterized (WPC, contralateral) sides in embWPC mice at E18. Control PMBSF and ALBSF territories were translated to the WPC side for direct comparison and labelled as “PMBSF” and “ALBSF”. Right, reslice of the yellow line in the left, representing responses elicited in PMBSF and ALBSF primordia by whisker pad stimulation (wps) or upper lip stimulations (uls). The white horizontal line delineates the boundaries between PMBSF and ALBSF, while the vertical dashed line divides the wps from the uls over time. Quantification of the PMBSF and ALBSF active fraction to wps or uls (n= 7 ctrl, n= 6 WPC, unpaired two-tailed Student *t*-test). **(C)** Left, cortical evoked responses (GCaMP6f) to whisker pad (magenta) and upper lip (cyan) stimulations in control (ipsilateral) and WPC (contralateral) sides in embWPC at P4. Right, reslice of the yellow line in the left, representing responses elicited in PMBSF or ALBSF by wps or uls. The white horizontal line delineates the boundaries between PMBSF and ALBSF, while the vertical dashed line divides the wps from the uls over time. Quantification of the PMBSF and ALBSF active fraction to wps and uls (n= 6 ctrl side, n= 6 WPC side, PMBSF active fraction, unpaired two-tailed Student *t*-test; ALBSF active fraction, Mann-Whitney *U*-test. ns, p= 0.06). **(D)** Quantification of the percentage of overlap responses between the PMBSF and ALBSF territories at E18 and P4 (n= 9 ctrl mice E18, n= 6 embWPC mice E18, n= 9 ctrl mice P4, n= 6 embWPC mice P4, Kruskal-Wallis test, ***p <0.001. Dunn’s multiple comparison test post-hoc analysis, ctrl E18 vs embwpc E18 ns, p= 0.37; embWPC E18 vs ctrl P4 ns, p= 0.77; embWPC E18 vs embWPC P4 ns, p= 0.14; ctrl P4 vs embWPC P4 ns, p > 0.99). **(E)** Schema illustrating the results found. Dashed lines delineate PMBSF and ALBSF putative evoked territories at E18. S1, primary somatosensory cortex; wps, whisker pad stimulation; uls, upper lip stimulation; GCaMP6f, calmodulin-based genetically encoded fluorescent calcium indicator 6–fast; WPC, whisker pad cauterized; contra, contralateral cortex to the whisker pad cauterized; ipsi, ipsilateral cortex to the whisker pad cauterized. Scale bars, 1000 μm. Bar graphs show the means ± SEM. ns, not significant. *p < 0.05, **p < 0.01, ***p < 0.001. See also Videos S3-S6.

Stimulations of either the whisker pad or the upper lip in the control side elicited responses from clusters of cortical neurons corresponding to the putative territories of the PMBSF or ALBSF, respectively (**Figure 2B**; **Video S3**). In contrast, whisker pad stimulations in the cauterized side (embWPC) triggered very reduced responses in the caudal-most part of the expected contralateral PMBSF (“PMBSF”), confirming the anatomical reduction of this territory. Remarkably, stimulations of the upper lip in the embWPC side elicited misplaced responses in the expected ALBSF (“ALBSF”) that notably expanded into the “PMBSF” (**Figure 2B**; **Video S4**), as compared to the control side. This ectopic “PMBSF” activation was maintained at P4 when the size of responses to upper lip stimulation were already refined in the control side (**Figures 2C-2E**; **Videos S5** and **S6**). Thus, the anatomical rescaling of barrel field sub-territories seen in the postnatal embWPC can be functionally detected prior to birth.

### Silencing prenatal thalamic synchronous activity does not prevent the area rescaling of barrel field sub-territories

Thalamic neurons display waves of prenatal spontaneous activity that are crucial for the development of cortical barrel field maps^10,21^. Therefore, we investigated whether the rescaling of barrel field territories in embWPC embryos could be due explained by changes in spontaneous activity of VPM neurons. Inputs from the whisker pad at E16 are carried by afferents from ventral portion of the PrV (vPrV), whereas inputs from the upper lip are carried by afferents from dorsal PrV (dPrV)^17–19^. While at this prenatal stage, vPrV afferents topographically target the dorsolateral VPM (the prospective wpVPM), dPrV afferents target not only the ventromedial part (the prospective ulVPM) but cover most part of the VPM^17,19^. Thus, while whisker pad and upper lip inputs are segregated at the brainstem nuclei, dPrV and vPrV axons terminals prenatally overlap within the prospective wpVPM territory^17,19^ (**Figures 3A** and **S6**). Remarkably, despite this axonal overlap, spontaneous activity patterns at E16 were sub-territory specific. We detected a notable distinction in the frequency of spontaneous activity waves between the prospective wpVPM and ulVPM regions. While wpVPM exhibited a high frequency of waves, the prospective ulVPM showed significantly reduced frequency (**Figures 3B** and **3C**; **Video S7**). In embWPC, we observed a significant change in this spontaneous activity pattern, with a striking increase in thalamic waves frequency within the prospective ulVPM, resembling the pattern observed in the control wpVPM primordium (**Figure 3B** and **3C**; **Video S8**). Thus, during prenatal development whisker input deprivation remaps spontaneous neuronal activity patterns in the future ulVPM territory. Next, we asked whether the embryonic increased activity in the prospective ulVPM might influence the functional and anatomical rescaling of the ALBSF territory. Thus, we tested this possibility and conducted embryonic whisker pad cauterization in a mouse in which the synchronous spontaneous activity of the thalamus is embryonically disrupted (referred to as embWPC-*Th^Kir^*)^21^. Briefly, we used a tamoxifen-dependent *Gbx2^CreERT^*^2^ mouse with a floxed line expressing the inward rectifier potassium channel 2.1 (*Kcnj2*) fused to the mCherry reporter in thalamic neurons (referred to as *Th^Kir^*). Our data revealed that thalamic and cortical territories in embWPC-*Th^Kir^* mice exhibited a spatial reorganization similar to that observed in embWPC mice. In both mice, the wpVPM and PMBSF territories virtually disappeared, and the ulVPM and ALBSF expanded as compared to the control (**Figure 3D**). Consistent to previous studies in the *Th^Kir^* mouse^21^, manipulating thalamic activity patterns in the embWPC mice (embWPC- *Th^Kir^*) resulted in the absence of thalamic waves (**Figures S7A** and **S7B**) and, the complete loss of barrels, evidenced by a total lack of TCA clustering, in both PMBSF and ALBSF territories (**Figures 3D** and **3E**). The rescaling of thalamocortical circuits was also detected in the embWPC-*Th^Kir^*mouse, but not in the *Th^Kir^* mouse alone, as evidenced by dye tracing studies (**Figures 3E** and **S7C**). In sum, our results demonstrate the patterns of spontaneous activity in prenatal VPM neurons are dispensable for intra-modal thalamic and cortical sub-territories rescaling to input deprivation.

**Figure 3.**
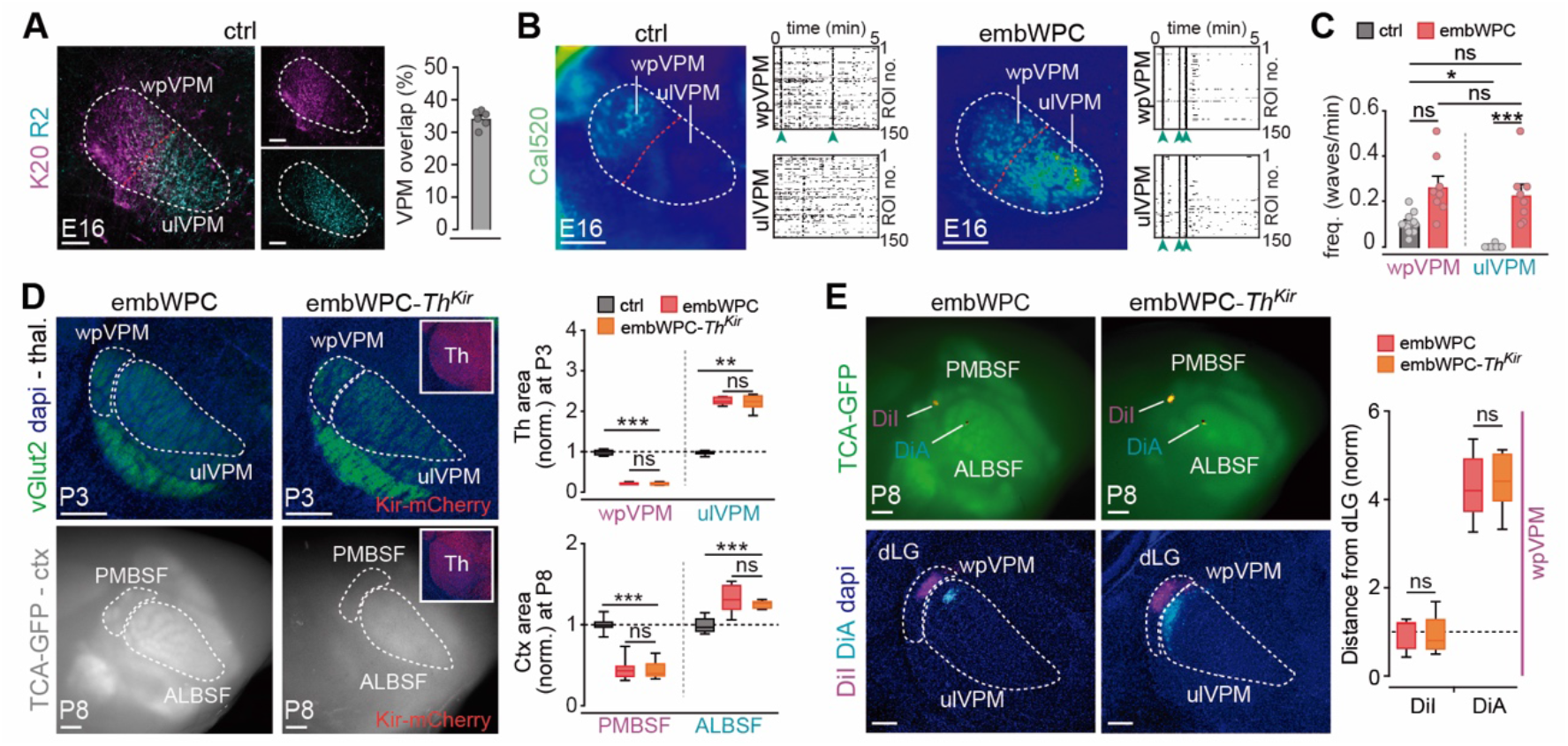
ALBSF rescaling is independent of thalamic patterned activity. (A) Coronal sections from the *Krox20-zsgreen::R2-mCherry* double transgenic mouse showing Krox20-labelled axons (magenta, vPrV) and R2-labelled axons (cyan, dPrV) at the VPM of the thalamus at E16. Quantification of the percentage of overlap of labelled axons at the thalamus (n= 6). **(B)** Maximal projection of *ex vivo* spontaneous calcium activity in the prospective VPM sub-regions (wpVPM and ulVPM) of the thalamus in acute slices at E16. Raster plots of 5 minutes. **(C)** Quantification of the frequency of waves activity in the VPM in both control and embWPC mice (n= 11 ctrl mice, n= 8 embWPC mice, Kruskal-Wallis test, ***p <0.001. Dunn’s multiple comparison test post- hoc analysis, prospective wpVPM ctrl vs prospective wpVPM embWPC ns, p= 0.14; prospective wpVPM ctrl vs prospective ulVPM embWPC ns, p= 0.32; prospective wpVPM embWPC vs prospective ulVPM embWPC ns, p >0.99). **(D)** Upper panels, coronal sections of vGlut2 staining (green) showing the size of the wpVPM and ulVPM in the thalamus of embWPC and embWPC-*Th^Kir^* mouse (Kir-mCherry positive) at P3. Lower panels, surface view of thalamocortical terminals (TCA-GFP+) in the cortical PMBSF and ALBSF in embWPC and embWPC-*Th^Kir^* mice at P8. Quantification of the data (Thalamus, n= 5 ctrl mice, n= 5 embWPC mice, n= 6 embWPC-*Th^Kir^* mice, wpVPM, Two-way ANOVA test: ***p <0.001. Tukey’s multiple comparison test post-hoc analysis, ns, p= 0.98; ulVPM, Kruskal-Wallis test, **p <0.01. Dunn’s multiple comparison test post- hoc analysis, ns, p >0.99. Cortex, n= 10 ctrl mice, n= 13 embWPC mice, n= 7 embWPC- *Th^Kir^* mice, PMBSF, Two-way ANOVA test: ***p <0.001. Tukey’s multiple comparison test post-hoc analysis, ns, p= 0.99; ALBSF, Two-way ANOVA test: ***p <0.001. Tukey’s multiple comparison test post-hoc analysis, ns, p= 0.52). **(E)** Upper panels, DiI and DiA crystal placements in the cortical PMBSF and ALBSF areas, respectively. Lower panels, backlabelled cells at the thalamus show shifted positions within the ulVPM thalamic area in both embWPC and embWPC-*Th^Kir^*mice at P8. Quantification of the position of backlabelled cells with respect to the distance to the dLG nucleus (n= 6 embWPC mice, n= 6 embWPC-*Th^Kir^* mice, DiI, Mann-Whitney *U*-test. ns, p= 0.7. DiA, unpaired two-tailed Student *t*-test; ns, 0.76). Scale bars, (A) 100 μm; (B) and (E, bottom) 200 μm; (D, top) 250 μm, (D, bottom) and (E, top) 500 μm. Boxplots show the medians with the interquartile range (box) and range (whiskers). Bar graphs show the means ± SEM. ns, not significant. *p < 0.05, **p < 0.01, ***p < 0.001. See also Figure S6 and S7; Videos S7 and S8.

### Upper lip input-receiving thalamic neurons adopt a similar transcriptional profile to mystacial input-receiving neurons

Cross-modal and intra-modal changes in the transcriptional programs of thalamic neurons have been shown to generate thalamocortical circuit re-organizations and cortical adaptations, as shown in early blind pups or in mice with an embryonic ablation of the VPM nucleus^10,23,24^. Therefore, we examined whether the anatomical and functional circuit expansion of the ALBSF in embWPC might be influenced by changes in the transcriptional program of VPM neurons. To assess this, we dissected tissue, at similar distances from the dorsal lateral geniculate (dLG) nucleus, from wpVPM and ulVPM in control and embWPC mice at P0 (**Figure 4A**) and compared their transcriptional profiles by bulk RNA-sequencing (RNA-seq). Principal component analysis (PCA) revealed that the transcriptomes from wpVPM and ulVPM cells in control mice clustered separately already at P0 (**Figure S8A**). Differential Expression Analysis (DEA) further revealed 377 Differentially Expressed Genes (DEGs) enriched in the wpVPM as opposed to 365 in the ulVPM cells. Among the DEGs enriched in each population, we found genes previously identified as being involved in somatosensory development such as *Epha3, Sox2, Hs6st2, Cdh8 or Rorα* enriched in wpVPM^24–28^ or *Rorβ, Slitrk6, Foxp2, Calb1 or Lhx2*^10,29–32^ enriched in the ulVPM (**Figures S8B** and **S8C**; **Table S1**).

**Figure 4.**
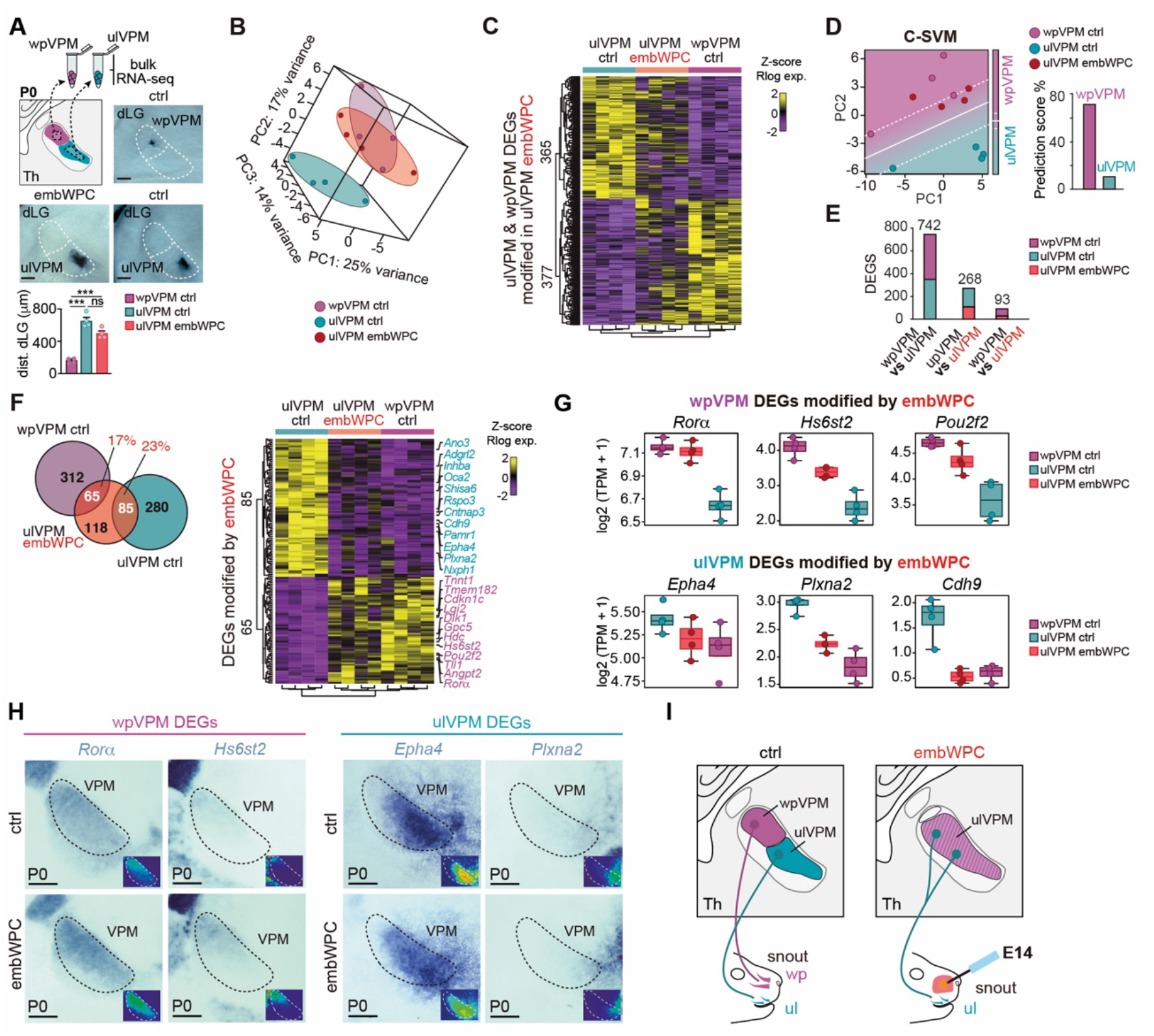
Upper-lip recipient thalamic neurons switch their transcriptional program to that of whisker-pad neurons. (A) Top left, schema representing the experimental paradigm used. Top right and bottom, coronal images showing the dissected thalamic territories for sequencing in control and embWPC slices. Quantification of the dissected area with respect to the distance to the dLG nucleus (n= 4 wpVPM ctrl slices from 4 mice, n= 4 ulVPM ctrl slices from 4 mice, and n= 4 ulWPC embWPC slices from 4 mice, One- way ANOVA test: ***p <0.001. Tukey’s multiple comparison test post-hoc analysis, ns, p > 0.05, ***p <0.001). **(B)** Principal Component Analysis (PCA) of wpVPM control (n= 4), ulVPM control (n= 4) and ulVPM whisker pad cauterization (embWPC, n= 4) at P0. **(C)** Heatmap of the normalized regularized logarithm (Rlog) Z-score of expression and unbiased clustering of region-specific DEGs showing their expression profiled in the ulVPM of the embWPC at P0. The color-code (yellow, high expression; purple, low expression) corresponds to the log2FC. **(D)** Left, C-support vector machine (C-SVM) analysis identifies the optimal demarcation plane between wpVPM (magenta dots) and ulVPM population (cyan dots). The ulVPM embWPC samples are identified as wpVPM control based on its gene expression pattern (red dots). Right, Prediction score value of ulVPM embWPC classified as wpVPM (70%) or ulVPM (30%) controls. **(E)** Number of Differentially Expressed Genes (DEGs) obtained in their respective differential expression analysis. **(F)** Left, Venn Diagram shows the number of genes modified by embWPC in ulVPM overlapped in every subset of region specific DEGs. Right, Heatmap of the normalized regularized logarithm (Rlog) Z-score of expression and unbiased clustering of region-specific DEGs whose expression was modified by embWPC in the ulVPM at P0. The color-code (yellow, high expression; purple, low expression) corresponds to the log2FC. **(G)** Boxplots showing TPM expression levels of selected wpVPM-specific and ulVPM-specific DEGs modified by embWPC in the ulVPM. Boxplots show the medians with the interquartile range (box) and range (whiskers). **(H)** Coronal sections showing by in situ hybridization the change in the pattern of expression of wpVPM and ulVPM DEGS in the embWPC mouse at P0 (n= 5 ctrl, n= 5 embWPC, for each probe). **(I)** Schema illustrating the results found. VPM, ventroposterior medial nucleus. Scale bars, 200 μm. See also Figures S8 and S9.

Then, we compared control versus embWPC transcriptional programs in the VPM. Remarkably, PCA analysis from ulVPM cells in embWPC mouse indicated that these neurons grouped more closely with control wpVPM than control ulVPM (**Figures 4B** and **S8D**). This transcriptional shift was appreciated when the expression pattern of the upper lip and whisker-pad DEGs was plotted in the ulVPM of the embWPC (**Figure 4C**). To investigate this effect, we trained a Support Vector Machine (C-SVM) classifier using region-specific genes and compared the results with the ulVPM-embWPC. This machine learning approach predicted that the cauterized model would be classified as the wpVPM control in 70% of cases, compared to 30% for the ulVPM control (**Figure 4D**). Additionally, DEA against the wpVPM control revealed a much lower number of differentially expressed genes (93 DEGs) compared to the ulVPM control (268 DEGs) (**Figure 4E**).

In total, 150 region-specific DEGs from ulVPM and wpVPM where significantly changed by the embWPC (**Figure 4F**). Namely, in the ulVPM of the embWPC mouse, we observed an upregulation of approximately 17% of the DEGs normally expressed in the control wpVPM and a downregulation of about 23% of control ulVPM DEGs, respectively (**Figures 4F** and **S8E**; **Table S2**). Notably, genes such as *Rorα, Epha4, Hs6st2, Cdh9,* or *Plxna2*, involved in axon guidance, thalamocortical mapping and wpVPM development^26,33^, exhibited a wpVPM-like pattern in the ulVPM of embWPC mice (**Figures 4G** and **S8F**), also corroborated by *in situ* hybridization (**Figures 4H** and **S8G**). Next, we performed a Gene Ontology (GO) functional enrichment analysis of the genes with expression changes in the ulVPM of the embWPC and identified 11 clusters of GO-Terms enriched in Biological Processes (BP) and Molecular Functions (MF) (**Figure S9**; **Table S3**). The top ranked cluster included 99 genes (46,8%) enriched in BP involved in dendrite development or synapse assembly (e.g. *Robo1, Flrt3*), and 12 genes (18,8%) enriched in MF involved in voltage-gated channel activity (e.g. *Kcnc2, Kcnab1*). In summary, the spatial rescaling of cortical ALBSF territory in embWPC correlates with transcriptomic changes observed in upper lip thalamic input neurons, which shift towards a molecular signature resembling that of whisker pad input neurons (**Figure 4I**).

### Increase of barrel size and functional spatial resolution of the cortical ALBSF map

We investigated whether upper lip-recipient thalamic neurons that acquire a whisker pad transcriptional profile during perinatal development might develop additional PMBSF characteristics later in life. At P8, PMBSF and ALBSF cortical areas differ in the average size of their barrels and the extent of clustering of thalamocortical axons. Remarkably, we observed that, in addition to the overall increase in the cortical area occupied by the ALBSF, the average size of the cortical individual barrels in the ALBSF of the embWPC was significantly larger compared to the ALBSF control (**Figure 5A**). This change appears to follow a gradient, with the largest ALBSF barrels located nearest to the remaining PMBSF. Furthermore, the cortical map of the expanded ALBSF gained definition and contrast as thalamocortical terminals appeared more clustered within each barrel, and less innervating the barrel septa, resembling the control PMBSF (**Figures 5B-5D**). These differences in barrel size and contrast in the ALBSF map of the embWPC where not detected in P0WPC mice (**Figures 5A** and **5B**).

**Figure 5.**
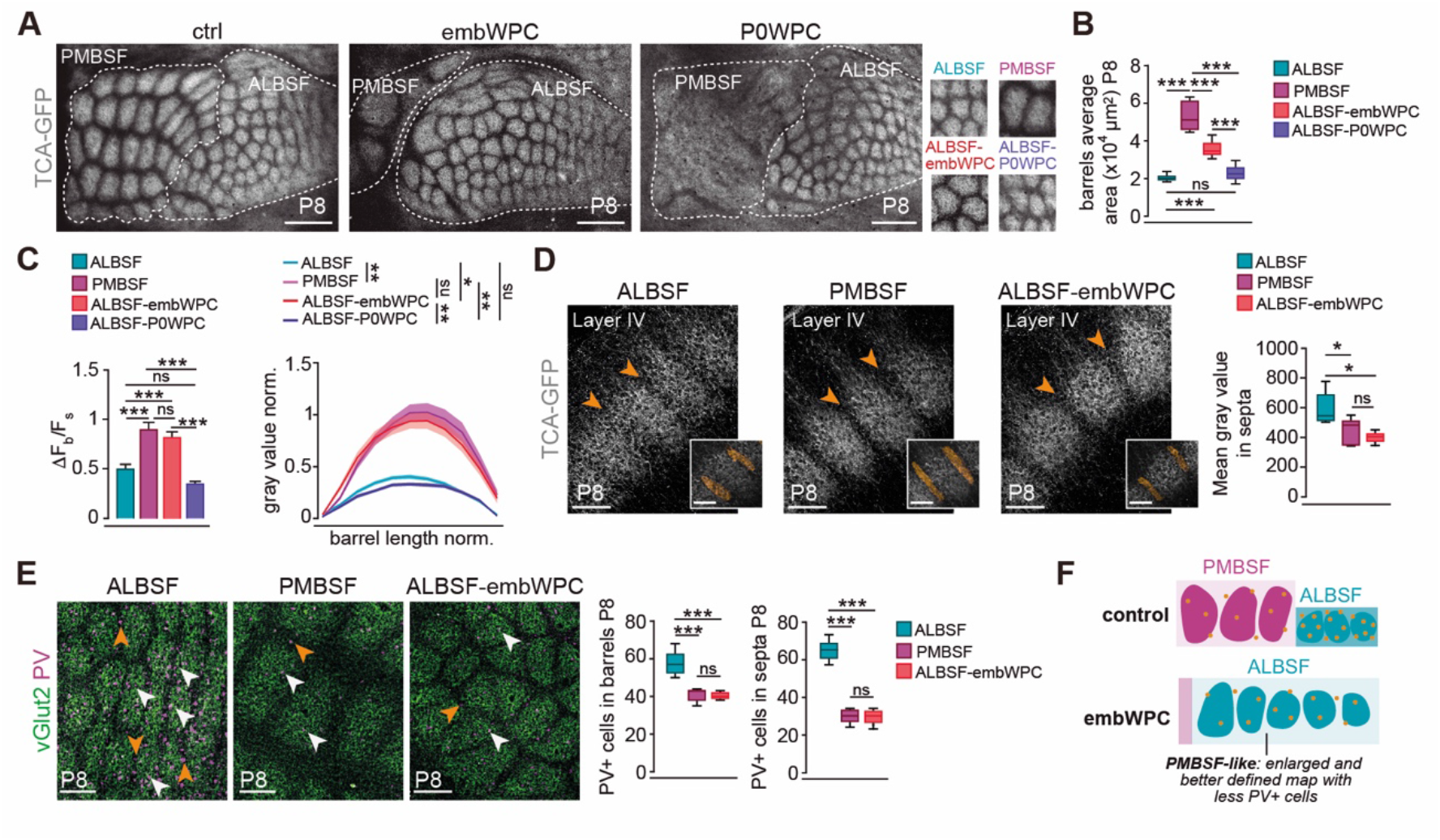
ALBSF barrels acquire PMBSF cortical features. (A) Left, cortical flattened tangential sections showing thalamocortical terminals (TCA-GFP+) in the PMBSF and ALBSF in control, embWPC and P0WPC mice at P8. Right, insets show in detail the clustering of thalamocortical axons in each condition. **(B)** Quantification of the data (barrel average area of 34 barrels, n= 7 ctrl mice PMBSF, n= 9 ctrl mice ALBSF, n= 9 embWPC mice ALBSF, n= 5 P0WPC mice ALBSF, Two-way ANOVA test: ***p <0.001. Tukey’s multiple comparison test post-hoc analysis, ns, p= 0.88). **(C)** Left, quantification of the barrel/septum fluorescence ratio (ι1Fb/Fs) (n= 12 ctrl mice PMBSF, n= 13 ctrl mice ALBSF, n= 13 embWPC mice ALBSF, n= 5 P0WPC mice ALBSF, Two-way ANOVA test: ***p <0.001. Tukey’s multiple comparison test post-hoc analysis, PMBSF-ctrl vs ALBSF- embWPC ns, p= 0.80; ALBSF-ctrl vs ALBSF-P0WPC ns, p= 0.35). Right, quantification of the gray value fluorescence intensity in a barrel (n= 66 ctrl PMBSF barrels, n= 65 ctrl ALBSF barrels, n= 42 embWPC ALBSF barrels, n= 25 P0WPC ALBSF barrels, One-way ANOVA test: ***p <0.001. Tukey’s multiple comparison test post-hoc analysis, ns, p= 0.99). **(D)** Coronal sections showing thalamocortical terminals in layer IV in the ALBSF and PMBSF of control, and ALBSF of embWPC mice. Insets showing the septa volume analyzed. Imaris quantification of the EGFP expression from TCA in septa of three barrels per animal (orange arrowheads) (n= 5 ctrl mice ALBSF, n= 5 ctrl mice PMBSF, n= 5 embWPC mice ALBSF, One-way ANOVA test: *p <0.05. Tukey’s multiple comparison test post-hoc analysis, ns, p= 0.75). **(E)** Cortical flattened tangential sections showing thalamocortical terminals (vGlut2+) and parvalbumin (PV) expression in barrel core (white arrowheads) and septa (orange arrowheads) in the ALBSF and PMBSF of control, and ALBSF of embWPC mice at P8. Quantification of PV+ cells in 4 barrels and each septa (n= 5 ctrl mice ALBSF, n= 5 ctrl mice PMBSF, n= 5 embWPC mice ALBSF, One-way ANOVA test: ***p <0.001. Tukey’s multiple comparison test post-hoc analysis, ns, p >0.05). **(F)** Schema illustrating the results found. Scale bars, (A) 500 μm, (D) 100 μm, (D, insets) 80 μm and (E) 200 μm. Boxplots show the medians with the interquartile range (box) and range (whiskers). Bar graphs show the means ± SEM. ns, not significant. *p < 0.05, **p < 0.01, ***p < 0.001.

Since thalamocortical input is a primary regulator of region-specific cortical features, we investigated whether other cortical characteristics, besides barrel size and thalamocortical clustering, had also changed in the ALBSF cortex of the embWPC. The postnatal PMBSF is known to contain fewer parvalbumin (PV) expressing interneurons in layer IV compared to the ALBSF^34^. Like in the PMBSF, we found significantly fewer PV-positive interneurons both in barrels and septa in layer IV of the ALBSF in the embWPC as compared to the ALBSF control (**Figure 5E**). Hence, our findings demonstrate that, irrespective of receiving input from upper lip peripheral receptors, the VPM territory in embWPC adopts a molecular identity akin to that of a territory receiving input from mystacial whiskers. Consequently, it gains specific features of that circuit, including increased barrel size, better anatomical map definition, and fewer layer IV PV interneurons (**Figure 5F**).

Next, we tested whether these anatomical reorganizations might have a functional correlate, as for example an enhanced topographical spatial resolution to upper lip stimulation. Thus, we carried out multi-point stimulations within the whisker pad or upper lip on both control and embWPC mice at P4 (**Figure 6A**). On the control mice, cortical evoked responses confirmed a topographic organization of both PMBSF and ALBSF responses; however, while the PMBSF responses exhibited a well-defined and precise organization with virtually non-overlapping responses, those of the ALBSF were broader and less refined in comparison (**Figure 6B**; **Video S9**). Thus, beyond their anatomical idiosyncrasies, cortical barrel sub-fields also have a distinct degree of functional map definition as they emerge during perinatal development. Interestingly, stimulations from distinct locations of the upper lip of embWPC mice resulted in significantly less overlapping responses within the ALBSF (**Figure 6B**; **Video S10**), suggesting an enhanced spatial discrimination in the ALBSF of embWPC akin to that seen in the control PMBSF map.

**Figure 6.**
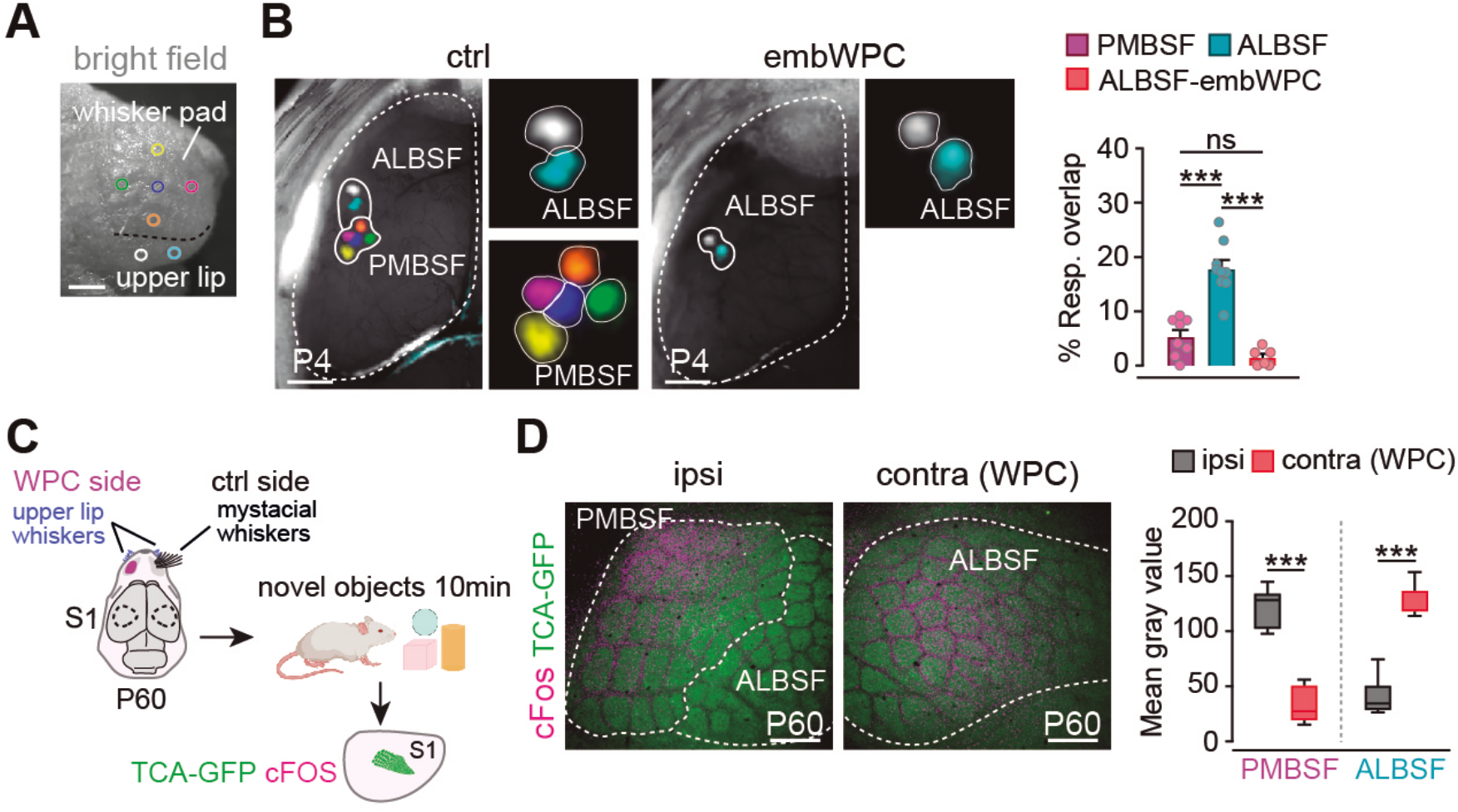
Enhance ALBSF functional map in the postnatal embWPC mice. (A) Schema representing the experimental paradigm. **(B)** Left, cortical evoked responses (GCaMP6f) to five distinct whisker pad stimulations and two distinct upper lip stimulations in control and embWPC mice at P4. Dashed lines indicate the territory covered by elicited responses within the PMBSF and ALBSF. Right, quantification of the percentage of overlap between two distinct PMBSF and two distinct ALBSF responses at P4 (n= 8 ctrl mice for PMBSF responses, n= 9 ctrl mice for ALBSF responses, n= 6 embWPC mice for ALBSF responses, Kruskal-Wallis test, ***p <0.001. Dunn’s multiple comparison test post-hoc analysis, ns, p= 0.18). **(C)** Schema representing the experimental paradigm. **(D)** Cortical flattened tangential sections showing thalamocortical terminals (TCA-GFP mouse) and cFOS expression in cortical ipsilateral (control) and contralateral (WPC) sides of P60 mice. Quantification of the mean gray value of cFOS expression in PMBSF and ALBSF in both sides (n= 8 embWPC mice ipsi PMBSF, n= 8 embWPC mice ipsi ALBSF, n= 8 embWPC mice contra (WPC) PMBSF, n= 8 embWPC mice contra (WPC) ALBSF, PMBSF, unpaired two-tailed Student *t*-test; ***p <0.001; ALBSF, Mann-Whitney *U*-test). Scale bars, (A) 1000 μm and (D) 500 μm. Bar graphs show the means ± SEM. ns, not significant. ***p < 0.001. See also Videos S9 and S10.

Finally, we investigated whether the rescaled ALBSF with enhanced resolution of in the embWPC mice takes on PMBSF functions during somatosensory behavior in adulthood. To do this, we conducted cFOS staining to label stimulated neurons in the ipsilateral and contralateral barrel sub-fields of embWPC mice after 10 minutes of exploring an enriched environment at P60 (**Figure 6C**). Our immunostaining for cFOS revealed that, whereas in the ipsilateral (control) side the ALBSF was barely activated as compared to the PMBSF, the ALBSF was the predominantly activated region in the contralateral (cauterized) side after exploration. These results strongly suggest that the remapped ALBSF neurons of the embWPC mouse play a prominent role in vibrissae- driven sensory behavior in the adult.

## DISCUSSION

Our study underscores the existence of a prenatal temporal window during mouse brain development that allows for the tuning of the size and definition of somatosensory cortical representations. Before cortical barrel fields are anatomically apparent, the thalamic VPM sub-territories exhibit specific transcriptional signatures at birth. Our results suggest that these signatures contribute to the distinct intra-modal anatomical and functional circuitry features specific to PMBSF and ALBSF cortical sub-territories.

Furthermore, our findings suggest that modulations in these transcription programs might enable intra-modal plasticity changes independently of the type of peripheral receptors. In mice lacking the mystacial whiskers, VPM neurons receiving upper lip whisker input can shift their transcriptional features toward a whisker pad-like program, reconfiguring thalamocortical circuits and cortical features to resemble a PMBSF. These changes increase barrel size and functional spatial resolution, thereby enhancing spatial discrimination in regions such as the ALBSF, where cortical representations were otherwise less defined.

The developmental relationship between sensory receptors and somatosensory representations has been thoroughly examined in prior work involving mice and rats. Embryonic ablation of the forelimb in rats resulted in the expansion of S1 representation of the hindlimb without altering the sensory receptors of the spared peripheral region^35^. Moreover, deprivation of a single whisker or a row of whiskers during the first postnatal week, leads to the loss of the corresponding cortical barrels and enlargement of the surrounding barrels^36–41^. And remarkably, rewiring of ipsilateral trigeminal input from mystacial whiskers to the ipsilateral barrel field generates an ectopic barrel field within the normal barrel map^42^. Collectively, these studies highlight the sensitivity of somatosensory regions and barrel maps to manipulations in peripheral or sensory input but also questioned a direct correlation between the size of cortical barrels and their corresponding peripheral sensory receptors. Our results in the embWPC model define a developmental window for an intra-modal barrel field reorganization, showing that only embryonic, and not postnatal, ablation of the whisker pad induces the ALBSF anatomical and functional enlargement. This reorganization capability is therefore temporally constrained during prenatal development and anticipates the closure of the critical time window for somatosensory circuits^38,43^.

Thalamic spontaneous activity is key for the emergence of cortical somatosensory maps, as evidenced by the fact that suppressing this activity leads to mice that do not develop barrels despite having normal whiskers^21^. Moreover, in early blind pups, changes in activity-dependent gene regulation in VPM neurons correlate to changes in barrel size^10^. Using embWPC, we could show that the enlargement of the cortical ALBSF area is guided by the thalamus in an activity-independent manner. Indeed, in embWPC-*Th^Kir^*mice, which lack synchronous thalamic activity, the rescaling of the ALBSF still occurs. On the other hand, similarly to what occurs cross-modally in early blind pups between the dLG and the VPM^10^, we observed a striking intramodal reorganization of the frequency and pattern of thalamic waves between VPM sub-fields in embWPC mice. The ulVPM significantly increased the frequency of thalamic waves compared to control mice. This change in activity pattern might serve as a mechanism to enhance ulVPM barrel size and definition and, in turn, to map these activity-dependent features onto the cortical ALBSF during development. This hypothesis could not be directly tested, as thalamocortical axons in the embWPC-*Th^Kir^* mice fail to cluster in layer IV in both PMBSF and ALBSF, as previously described in *Th^Kir^* mice^21^.

Remarkably, our study revealed that thalamic neurons can shift their transcriptional profile intra-modally. We found that the transcriptional profile of ulVPM neurons in embWPC mice mirrors that of the wpVPM neurons in control mice. Crucial genes for thalamocortical barrel field development, such as *Rorα* or *Epha4*^26,33^, accordingly altered their expression pattern in the ulVPM neurons innervating the ALBSF. For instance, we observed an upregulation of *Rorα* in this region, suggesting a potential role in increasing the density and clustering of axonal terminals in layer IV in the ALBSF of the embWPC mice. An activity-dependent upregulation of *Rorβ*, a related orphan receptor, in VPM neurons has been shown to mediate an increased clustering of thalamic axons in S1^10^. ulVPM neurons adopting a region-specific wpVPM transcriptional profile might serve as an intrinsic mechanism to switch a PMBSF developmental program, ensuring an enhanced anatomical and functional definition of thalamocortical axons in the ALBSF cortical barrel map. Additionally, we identified alterations in genes directly associated with neuronal activity^44,45^, including ion channels, which may contribute to or reflect the modified pattern of thalamic activity observed in the embWPC model. Yet, it remains to be determined whether the acquisition of wpVPM-like molecular identity by ulVPM neurons is dependent on thalamic spontaneous activity.

Large barrels in the PMBSF correspond to long mystacial whiskers^12,13^, whereas ALBSF barrels relate to the short whiskers on the upper lip of the snout and are comparatively smaller, less defined and densely packed^9,46,47^. Moreover, although the average volume of a single barrel in the ALBSF is smaller than in the PMBSF, the numerical density of parvalbumin (PV)-positive interneurons is higher in ALBSF than in the PMBSF^34^. Somata of PV neurons in the barrels and septa of the ALBSF are reported to receive more vesicular glutamate transporter Type 2–labeled (vGlut2) boutons than those in the PMBSF, suggesting the presence of more potent feedforward inhibitory circuits in the ALBSF^34^. Behaviorally, mystacial vibrissae hold significant behavioral relevance for the animal, being the first ones used for search of objects and gaps, and to discern diverse tactile features such as object position, shape, or texture^48^. Although the function of upper lip shorter whiskers remains largely unexplored, they appear to primarily aid in object discrimination when the snout is near the object^8,49–51^. Given these anatomical and functional disparities, the PMBSF and ALBSF represent two distinct somatosensory subsystems. Consequently, we initially expected minimal reorganization of the ALBSF upon deprivation of the whiskers corresponding to the PMBSF. However, our data from the embWPC mice revealed a notable anatomical reorganization within the ALBSF, leading to the formation of large and well-defined barrels and an increased density of PV interneurons at the postnatal life. Additionally, our *in vivo* functional data revealed that this reorganization might contribute to enhance functional resolution of the ALBSF map to point-to-point peripheral stimulations.

Altogether, these findings highlight a decoupling of mechanisms governing the type, size, and density of sensory peripheral receptors from those influencing the size and definition of cortical representations that might involve adopting subcortical- and subregion-specific transcriptional programs. Specifically, we observed that the prenatal brain possesses plasticity mechanisms to develop a barrel field with anatomical and functional features akin to the PMBSF, regardless of the type of peripheral receptor. Therefore, our findings indicate that the brain may still construct a barrel map capable of processing intricate tactile functions for the animal, even in scenarios involving the availability of only small receptors with subsidiary functions.

### Methods Mouse strains

All animal experiments were conducted in accordance with the Committee on Animal Research at the University Miguel Hernández that approved all the animal procedures, which were carried out in compliance with Spanish and European Union regulations. Similar numbers of male and female mice were used interchangeably. No sex-related differences were observed in the measurements throughout the study. Mice were maintained in pathogen-free facilities under standard housing conditions with continuous access to food and water on a 12h light-dark cycle. The number of animals used in each experiment is noted in the figure legends.

All mouse transgenic lines in this study were maintained on an ICR/CD-1 genetic background and genotyped by PCR. The *TCA-GFP Tg*, in which the TCAs are labelled with GFP, the *R26^Kir^*^2^*^.1-mCherry^*, mouse lines were previously described^10,14,21^. The Cre-dependent mouse line, *R26^GCaMP6f^* was obtained from Jackson Laboratories (Stock number 024105) and crossed with an *Emx1^Cre/+^*transgenic mouse^52^ to conditionally express the fast calcium indicator GCaMP6f in glutamatergic cortical neurons (*Emx1^GCaMP6f^*)^21,22^. The *R26^Kir2.1-mCherry^* mice were crossed with an inducible *Cre^ERT2^* mouse line driven by *Gbx2*, an early specific thalamic promoter (*Gbx2^CreERT2/+^*)^53^. Double mutants are referred as *Th^Kir^* and triple mutants as *TCA-GFP-Th^Kir^*^21^. Tamoxifen induction of Cre recombinase in the double/triple mutant embryos was performed by gavage administration of tamoxifen (5 mg dissolved in corn oil, Sigma) at E10 to specifically target all primary sensory thalamic nuclei. Tamoxifen administration in pregnant mice produces non-desirable side effects such as delivery problems and decrease survival of newborn pups^54^. To increase the survival rate of young pups, we administered 125 mg/Kg of progesterone (DEPO-PROGEVERA®) intraperitoneally at E14 and implemented C-section procedure at E19. Pups were then placed with a foster mother. In all cases, the Cre^ERT2^-negative littermates were used as controls of the experimental condition. The *r2^mCherry^* ^19^, *Krox20^Cre^* ^55^ and *R26RZsGreen* (Jackson Laboratories, Stock number 007906)^56^ lines, were as described. We crossed *Krox20^Cre^* with *R26RZsGreen* to generate double transgenic *Krox20^ZsGreen^* mice and we also generated *Krox20^ZsGreen^::r2^mCherry^*triple transgenic mice.

### Histology

For *in situ* hybridization and immunohistochemistry at postnatal stages, mice were perfused with 4% paraformaldehyde (PFA) in PBS (0.01 M), and their brains were subsequently removed and post-fixed in the same fixative overnight. For immunohistochemistry of embryonic tissue, the brains were dissected out and immediately fixed in 4% PFA overnight. For Nissl staining, a microtome (MICROM) was used to cut paraffin slices of 5 μm. Next, the sections were stained in 0.5% cresyl violet (Sigma) solution for 15-25 min and then rinsed quickly in distilled water. After decolorization in 70% ethyl alcohol for few seconds, the sections were dehydrated in 95%, 100% ethyl alcohol for 2 to 3 min, cleared in xylen (Sigma) for 2 min and mounted with Eukitt (Merk). Cytochrome oxidase staining was performed to label the PrV territories. For cytochrome Oxidase (CytOx) staining, 80 μm vibratome coronal sections were incubated overnight at 37 °C in a CytOx solution: 0.03% cytochrome c (Sigma C2506), 0.05% 3-3′ diaminobenzidine tetrahydrochloride hydrate (DAB, Sigma D5637) and 4% sucrose in PBS. For tangential sections, cortical hemispheres were flattened and cryoprotected through steps of 10%, 20% and 30% of sucrose in PBS. Then, a cryotome (MICROM) was used to cut at 80 μm tangential sections. Immunohistochemistry was performed on 80 μm vibratome or cryotome brain sections (coronal and tangential), which were first incubated for 1h at room temperature in a blocking solution containing 1% BSA (Sigma) and 0.25% Triton X-100 (Sigma) in PBS. Afterwards, the slices were incubated overnight at 4 °C with the following primary antibodies: guinea pig anti-vGlut2 (1:10000, Synaptic Systems, 135404), chicken anti-GFP (1:3000; Aves Labs, GFP-1020), rat anti-RFP (1:1000, Chromotek, 5F8), rabbit anti-cFos (1:500, Synaptic Systems, 226003), rabbit anti-PV (1:5000, Swant PV27), and rabbit anti-RFP (1:1000, Rockland, 600-401-379). Sections were then rinsed in PBS and incubated for 2h at room temperature with secondary antibodies: Alexa488 donkey anti-guinea pig (1:500, ThermoFisher, A11073), Alexa488 goat anti-chicken (1:500, ThermoFisher, A11039), Alexa594 donkey anti-rat (1:500, ThermoFisher, A21209) and Alexa goat 568 anti-rabbit (1:500, Invitrogen, A11011). Counterstaining was performed using the fluorescent nuclear dye 4′,6-diamidino-2-phenylindole (DAPI) (Sigma). In situ hybridization was performed on 60 μm vibratome sections using digoxigenin-labelled antisense probe for *Rorα*, *Epha4*, *Pou2f2*, *Hs6st2*, *Plxna2* and *Cdh9*. Hybridization was carried out overnight at 65 °C, and after hybridization, the sections were washed and incubated overnight at 4 °C with an alkaline phosphatase-conjugated anti-digoxigenin antibody (1:2500-1:4000, Roche). To visualize the RNA-probe binding, colorimetric reaction was performed for 1-2 days at room temperature in a solution containing NBT (nitro-blue tetrazolium chloride, Roche) and BCIP (5-bromo-4-chloro-30-indoly phosphate p-toluidine salt, Roche). After development, the sections were washed and mounted in Glycerol Jelly (Merck Millipore). Images were acquired with a Leica DFC550 camera into a Leica DM5000B microscope, Leica K5 camera into Leica DMi8 microscope or with an Axioscan Z1 widefield microscope (Zeiss).

### Immunolabeling-enabled three-dimensional imaging of solvent-cleared organ (iDISCO+)

Whole mount for the iDISCO+ protocol was conducted following the previously described methods^57,58^. After perfusing mice with 4% PFA, heads (without any dissection) were dehydrated using a series of methanol concentrations (50, 80, 100, and 100%) and subsequently incubated overnight in 6% H2O2 in methanol to bleach the samples. Following this, heads were blocked with PBS-GT (0.5% Triton X-100, 0.2% gelatin) for 4 days. For the clearing process, heads underwent dehydration in methanol (20, 40, 60, and 80%) at room temperature on a rotating shaker. Specimens were then immersed twice in 100% methanol for 1 hour and treated overnight in 1/3 volumes of 100% dichloromethane (DCM; Sigma-Aldrich; 270997). The subsequent day, heads were incubated in 100% DCM for 30 minutes. Finally, samples were cleared in 100% dibenzyl ether (DBE; Sigma-Aldrich; 108014) until they became translucent.

### Ultramicroscopy and Image Processing

3D imaging was primarily performed with an ultramicroscope I (LaVision BioTec) using ImspectorPro software (LaVision BioTec). The light sheet was generated by a laser (wavelength 488 nm, LaVision BioTec). A binocular stereomicroscope (MXV10, Olympus) with a 23x objective (MVPLAPO, Olympus) was used at different magnifications (1.25x). Samples were placed in an imaging reservoir made of 100% quartz (LaVision BioTec) filled with ethyl cinnamate and illuminated from the side by the laser light.

Images were generated using Imaris x64 software (version 9.3.1, Bitplane). Stack images were first converted to Imaris files (.ims) using ImarisFileConverter. The whisker pad and upper lip areas reconstruction were generated by creating a mask around each one using the “surface tool” and they were pseudo-coloured (whisker pad area in magenta and upper lip area in cyan). Each individual follicle was isolated also manually using the same tool, selecting nine follicles arbitrarily at three different points from medial to lateral. The septa volume reconstruction from coronal sections of 80 μm was generated by creating masks around each barrel using the “surface tool” and they were pseudo-coloured (orange).

### *In utero* and postnatal unilateral whisker pad cauterization

Embryonic unilateral whisker pad cauterization (embWPC) was performed at E14 as described previously for enucleation^10^. Dams were deeply anesthetized with isoflurane and the uterine horns were exposed through a midline laparotomy. After cauterizing right whisker pad side in half of the litter, the embryos were returned to the abdominal cavity, the surgical incision was closed, and the embryos were allowed to develop until either E18 or postnatal stages. Postnatal unilateral whisker pad cauterization (P0WPC) was performed on P0 pups. Animals were deeply anaesthetized on ice. The right whisker pad was cauterized under the loupe to specifically burn the principal whiskers follicles. Pups were then warmed up to 37°C on a heating pad, before being returned to the mother.

### Measurement of brain areas and data analysis

ImageJ software was used to measure the size of individual barrels, barrel field territories in the cortex, thalamus and brainstem, as well as the snout areas in slices. For barrel field territories and snout areas data were normalized. Each barrel field or snout area from a given experimental condition was normalized to the corresponding barrel field or snout mean area in the control, which was considered as 1. For the snout measurements, the skin of the snout was flattened and post-fixed in 4% PFA. Paraffin slices of 5 μm were obtained to quantify the number of follicles and the upper lip snout area using Nissl staining. We measure two slices per animal, one medial and one more lateral, and calculated the average. TCA-GFP mouse was used for the quantifications of the barrel field definition in flattened tangential sections, barrel field areas *in toto* and thalamic territories in slices. To quantify the size of cortical areas *in toto*, TCA-GFP (control, embWPC, P0WPC and embWPC-*Th^Kir^*) mice were perfused and directly processed to obtain images under the stereo fluorescent microscope (Leica MZ10 F). Coronal serial slices of 80 μm were obtained from TCA-GFP brains, and thalamic barrel field territories were immunolabel with GFP and vGlut2 in order to better detect the areas. For ι1Fb/Fs quantifications, we used ImageJ to measure the gray value of EGFP labelling in barrels of flattened tangential sections. ι1Fb/Fs was calculated using the maximum value of the baseline signal average as Fs in single barrels. To measure the gray value, we used a 20-width segmented line, covering 5 barrels per condition, and calculated the average. To compare the barrel profile for the different conditions, we normalized each barreĺs gray values to the lowest number (septa), which was considered as 0. Next, we took 11 gray value data points per barrel to normalize the length for all the conditions. To quantify the mean gray value in septa, coronal serial slices of 80 μm were obtained from TCA-GFP brains, and barrels from PMBSF and ALBSF (control and embWPC) were immunolabel with GFP to measure the fluorescence of three barrels septa per condition using Imaris x64 software, and calculated the average. For PV+ cells quantifications in PMBSF and ALBSF barrels and septa, we counted the number of PV+ cells within four barrels and their septa for each condition in flattened tangential sections of 80 μm immunolabel with PV and vGlut2. We quantified cFos expression within PMBSF and ALBSF layer IV. The region of interest (ROI) was a square of 600x1000 μm on each territory measuring the mean gray value.

### Dye-tracing studies

For axonal tracing, animals were perfused with 4% PFA in PBS. Heads were post-fixed overnight to trace innervation from the snout to the trigeminal nucleus, while brains were dissected and post-fixed overnight to trace thalamocortical axons. Small DiI (1,1′- dioctadecyl 3,3,3′,3′-tetramethylindocarbocyanine perchlorate; Invitrogen), DiA (4-[4- (dihexadecylamino) styryl]-N-methylpyridinium iodide; Invitrogen) and DiD (1,1’- Dioctadecyl-3,3,3’,3’-Tetramethylindodicarbocyanine, 4-Chlorobenzenesulfonate Salt; Invitrogen) crystals were inserted under a stereo fluorescence microscope (MZ10 F, Leica) into the PMBSF, ALBSF, the whisker pad and upper lip. The dye was allowed to diffuse at 37 °C in PFA solution for 4 to 8 weeks to trace the innervation from the snout to the PrV. To trace the thalamocortical pathway, small DiI and DiA crystals were inserted into distinct PMBSF barrels (C2, C3, D3 and C4) and DiI, DiA and DiD crystals were placed into ALBSF territory. To reveal the PBMSF and ALBSF, a TCA-GFP transgenic specific mouse line was used. The dye was allowed to diffuse at 37 °C in PFA solution for 3 weeks. Vibratome sections (80 μm thick) were obtained and counterstained with the fluorescent nuclear dye DAPI (Sigma-Aldrich). Three sections per animal were imaged using a Leica K5 camera into Leica DMi8 microscope. Image analysis was performed on ImageJ. To quantify the translocation of the backlabeling in the VPM, the distance from the dorsolateral geniculate (dLG) nucleus border was measured to the center of the backlabeled cells in the VPM. The distance from the dLG nucleus border to the separation between wpVPM and ulVPM was determined following the barreloids rows in the VPM of ctrl mice.

### Measurement of PrV axons overlap in VPM and data analysis

To label rhombomere 3-derived vPrV neurons, we used the *Krox20^Cre^* line^55^ to drive the Rosa-Zsgreen reporter (*R26RZsGreen*)^59^, whereas to label rhombomere 2-derived dPrV neurons, we used the *r2^mCherry^* reporter line^19^. Brains from *Krox20^Cre^*; *R26RZsGreen*; *r2^mCherry^* animals, in which vPrV and dPrV projections are labeled simultaneously, were collected at embryonic stages (E16.5, E18.5) and postnatal stages (P0, P4, P8) for analysis. Embryonic brains were directly post-fixed in 4% PFA in PBS, while postnatal brains were perfused before post-fixation in the same fixative. Brains were cut on a Vibratome (Leica) at 60 μm thickness throughout the VPM and PrV. To enhance the r2::mCherry signal, the sections were stained with a rabbit anti-RFP primary antibody (1:1000, Rockland, 600-401-379) coupled to an anti-rabbit Alexa 568 secondary antibody (1:500, Invitrogen, A11011), while the endogenous Zsgreen fluorescence was not amplified. At least five sections per animal were imaged using Axioscan Z1 widefield microscope (Zeiss), using a 10x (NA=0.45) plan-APOCHROMAT objective. Subsequent image analysis was conducted using ImageJ. To quantify the percentage of overlap of vPrV-dPrV axons in the VPM, wand tool connecting pixels of the same intensity level in a 15-pixel range, was used to automatically delineate the innervation area of Krox20+ and R2+ axons, generating the Regions of Interest (ROIs). Subsequently, the overlapping area and the total innervation area of both Krox20+ and R2+ axons were measured with the measure tool in ImageJ to calculate the percentage of overlap of PrV axons in the VPM. The same strategy was used for PrV dye tracings to measure the overlap between dPrV and vPrV axons in the VPM.

### *In vivo* mesoscale calcium imaging

As previously described^22^, embryos at E18 were extracted from the uterus and maintained at 35 °C. E18 pups were immobilized using soft clay. P4 mice underwent anesthesia with ice, followed by surgical removal of the scalp. A 3D-printed plastic holder was attached to the skull using cyanoacrylate adhesive and dental cement, then affixed to a ball-joint holder to stabilize the head. To maintain body temperature, pups were placed on a controlled temperature heating pad, ensuring a range of 32-34 °C. For recording calcium activity, we used a 16-bit CMOS camera (ORCA-Flash 4.0, Hamamatsu) coupled to a stereo microscope (Stereo Discovery V8, Zeiss), which offered 470 nm LED illumination. For *Emx1^GCaMP6f^*, images were acquired with a frame size of 1024x1024 pixels using a macro magnification of 1.6x at E18 and of 1.25x at P4, resulting in spatial resolution of 8.12 μm/pixel and 10.64 μm/pixel, respectively. Image frames were captured continuously at a rate of 3.33 frames per second (300 ms frame period) for *Emx1^GCaMP6f^* with an average of 3 movies was acquired per animal.

### *In vivo* mechanical stimulation of the whisker pad

We conducted somatosensory stimulations by touching the whisker pad and upper lip using a 0.16 g von Frey filament (TouchTest®, BIOSEB). Each animal was stimulated at least three times, with intervals of 5 minutes between each stimulus. Subsequently, we calculated the size of responses in the PMBSF and ALBSF elicited by whisker pad and upper lip stimulations.

### Analysis of the evoked activity *in vivo*

For the assignment of cortical territories, the perimeters of the PMBSF and ALBSF were determined using the cortical responses elicited by mechanical stimulation of five sites on the whisker pad, as previously described^21,22^ and two sites of the upper lip. At embryonic day (E)18, the perimeters were predicted by scaling down and superimposing the limits of the sensory territories observed in the TCA-GFP transgenic line^14^ at P2, as previously detailed^22^, and using the responses to whisker pad and upper lip stimulations as reference for PMBSF and ALBSF, respectively. Image analysis was performed with ImageJ. The mean fluorescence of five frames just prior to the stimulation established F0 (baseline fluorescence), and a ΔF/F0 time series was then generated from the raw data and subsequently transformed into 8-bit images before processing with a 2-pixel diameter Gaussian filter. A maximum intensity projection of all frames containing evoked responses was obtained. The boundaries of the response were defined using the wand tool, connecting pixels of the same intensity level within a 15-pixel range, and the area of each response was calculated using the measure tool in ImageJ. To measure the active fraction of PMBSF and ALBSF in embWPC mice at E18 and P4, the control side was used to delineate the PMBSF and ALBSF territories following stimulation of the whisker pad or upper lip, respectively. On the WPC side, these areas were superimposed using the mirror image from the control side. The PMBSF and ALBSF active fractions were determined by measuring all elicited responses within each territory and calculating the percentage of the PMBSF and ALBSF theoretical areas occupied by these responses following whisker pad or upper lip stimulations. To quantify the percentage of overlap between responses following whisker pad and upper lip stimulations (%PMBSF- ALBSF overlap) at E18 and P4, the overlap between all elicited responses following whisker pad and those following upper lip stimulations was calculated. To quantify the percentage of overlap within PMBSF or ALBSF (% Responses overlap) in ctrl and embWPC, the overlap for two different stimuli on the whisker pad or two different stimuli on the upper lip was calculated.

To visualize the evoked stimuli in different color profiles for the generation of supplementary movies, we utilized custom scripts developed in Matlab^TM^. These scripts were adapted from the image analysis suite WholeBrainDX (referenced in the Data and code availability section). For each movie, frames containing movement artifacts, spontaneous activity, and calcium activity deemed as stimulation byproducts were excluded. Baseline correction was performed using the built-in Matlab function ’msbackadj’ with a window size of 20 and a step size of 20. Subsequently, ι1F/F0 was computed using the median value of the corrected signal as F0. Movie segmentation and calcium event detection followed the method described^60^. Image segmentation and generation of a binary movie involved Gaussian smoothing with an 80 μm distance and a signal intensity threshold. The binary movie was color-labeled using ImageJ software and customized with a Gaussian filter. Subsequently, the original movie and the color-coded binary movie were merged using the ’Image Calculator’ function in ImageJ.

### Microdissection and RNA isolation for RNA-seq

To collect tissue from the wpVPM and ulVPM territories, ctrl and embWPC P0 pups were euthanized via decapitation, and their brains were dissected out under RNase-free conditions to prevent RNA degradation. The brains (five brains were pooled for each sample) were collected in ice-cold KREBS solution and sliced into 300 μm sections using a vibratome (VT1000S Leica). The wpVPM and ulVPM territories were rapidly microdissected under a stereo microscope. The bulk tissue was immediately transferred to lysis buffer of the Rneasy® Micro Kit (Qiagen, 74004) for total RNA extraction, following the manufacturer’s instructions. RNA quality was measured for all samples using an Agilent Bioanalyzer 2100 system, and only samples with RNA Integrity Number (RIN) > 8 were used for library construction.

### Library preparation and RNA sequencing

Library construction and sequencing were performed at Novogene Co. Ltd. Genomics core facility (Cambridge, UK). cDNA multiplex libraries were prepared using a custom Novogene NGS RNA Library Prep Set (PT042) kit. Briefly, mRNA was purified from total RNA using poly-T oligo-attached magnetic beads. After fragmentation, the first strand cDNA was synthesized using random hexamer primers followed by the second strand cDNA synthesis. The library was ready after end repair, A-tailing, adapter ligation, size selection, amplification, and purification.

The library was checked with Qubit and real-time PCR for quantification and bioanalyzer for size distribution detection. Libraries were pooled and sequenced in 2x150bp paired-end mode on a S4 flowcell in the Illumina Novaseq6000 platform. A minimum of 40 million reads were generated from each library.

### Bioinformatic analysis of the RNA-seq

RNA-seq analysis were performed as previously described^61^ with minor modifications: quality control of the raw data was performed with FastQC (v.0.11.9). RNA-seq reads were mapped to the Mouse genome (GRCm39) using STAR (v2.7.9a)^62^ and SAM/BAM files were further processed using SAMtools (v1.15). Aligned reads were counted and assigned to genes using Ensembl release 104 gene annotation and FeatureCounts, Subread (v2.0.1). Normalization of read counts and differential expression analyses were performed using DESeq2 (v1.32)^63^, Bioconductor (v3.15)^64^ in the R statistical computing and graphics platform (v4.2.2 “Innocent and Trusting”).

In the analysis of wpVPM and ulVPM datasets (control and embWPC samples) generated for this study, significantly Differentially Expressed Genes (DEGs) were identified using a simultaneous statistical significance threshold (Benjamini-Hochberg (BH) adjusted *P*-value < 0.1) and absolute log2 fold change (log2FC) > 0.14 by shrunken log2FC using the adaptive *T* prior Bayesian shrinkage estimator “apeglm” (**Tables S1** and **S2**)^65^. Hierarchical clustering analysis was performed using “Euclidean” distance and “Complete” clustering methods metrics to visualize significantly upregulated and down-regulated genes. A linear support-vector machine (SVM) model for classifying RNA-seq samples was developed using the e1071 package (v1.7-14). This model was based on the gene expression profiles of the top 500 most variable genes in ulVPM and wpVPM control samples. Subsequently, the C-SVM model was utilized to predict the classification of ulVPM embWPC samples based on their transcriptomic profiles.

Functional enrichment analyses were performed using clusterProfiler (v4.4.4)^66^ under org.Mm.eg.db package (v3.15) for better annotation data. All enriched terms were considered significant at adjusted P-values by ‘BH’ < 0.1, in the Gene Ontology (GO) Over-Representation Analysis. Enrichment results were further clustered and simplified using the simplifyEnrichment package (version 1.11.1)^67^ (**Table S3**).

### *Ex vivo* calcium imaging

At E16, embryos were retrieved from the dam’s uterus by cesarean section, their brains were rapidly dissected out and they were submerged in an ice-cold slicing solution containing (in mM): 2.5 KCl, 7 MgSO4, 0.5 CaCl2, 1 NaH2PO4, 26 Na2HCO3, 11 glucose and 228 sucrose. Coronal slices (350 μm thick) were obtained using a vibratome (VT1200S Leica) and they were left to recover for at least 30 minutes at room temperature in standard artificial cerebrospinal fluid (ACSF) containing (in mM): 119 NaCl, 5 KCl, 1.3 MgSO4, 2.4 CaCl2, 1 NaH2PO4, 26 Na2HCO3 and 11 glucose. All extracellular solutions were continuously bubbled with a 95% O2 and 5% CO2 gas mixture. Slices were loaded with the calcium indicator Cal520TM (AAT Bioquest) as previously described^10,21^, transferred to a submersion-type recording chamber and perfused with warm ACSF (32-34 °C) at a rate of 1.8 ml/min. Images were acquired with a digital charge-coupled device (CCD) camera (ORCA-R2 C10600-10B, Hamamatsu) coupled to an upright microscope (DM-LFSA, Leica) and using a 5x objective. For recordings of spontaneous calcium activity, frames were acquired with an exposure time of 150 ms, an interframe interval of 300 ms, a frame size of 672x512 pixel and a spatial resolution of 2.5 μm/pixel. With each slice, 1 to 5 epochs of 15 mins (3000 frames) were recorded.

### Analysis of fluorescence spontaneous activity

Analysis of spontaneous thalamic calcium activity was conducted using custom software developed in Matlab^TM^, which was adapted from the CalciumDX toolbox (indexed in the Data and code availability section). For each movie, VPM was delineated, and the two prospective areas (wpVPM and ulVPM) were subdivided into a grid of 6x6 pixels where each small square is a region of interest (ROI). Calcium activity events were detected based on the average calcium signal of each ROI over time, employing threshold-based algorithms modified from CalciumDX. To identify significant synchronous activity and discard ROI co-activation that resulting from random temporal coincidence of calcium events, we generated surrogated calcium events sequences for each experiment using Matlab^TM^. The alternative dataset was built by randomly shuffling the original temporal intervals between calcium transients in every ROI while preserving the spiking frequency and temporal structure of the calcium activity. Subsequently, the maximum value of co-activation from shuffled data was calculated. Through 1000 iterations, we established a synchronicity threshold as the 95^th^ percentile of the maximum values of co-activation obtained from the shuffled dataset. In each experiment, only activity super passing the synchronicity threshold was used for calculations and visualization. The onset of a synchronic event was defined as the frame in which co-activation overpass the threshold, while the end was determined as the frame in which co-activation reached 25% of peak synchronicity.

### Quantification and statistical analysis

Data were analysed using Prism 9 (GraphPad). A Kolmogorov-Smirnov normality test was conducted on all datasets. For independent data that conforming to a normal distribution, an unpaired two-tailed Student’s *t*-test was employed to compare two groups. In cases where independent data that did not follow a normal distribution, a Mann-Whitney *U*-Test two-tailed test was used for comparison. For analyses involving more than two groups and one factor, one-way ANOVA was applied, followed by a Tukey *post hoc* analysis when data exhibited a normal distribution. For more than two groups and two factors, two-way ANOVA was conducted, followed by a Tukey *post hoc* analysis for normally distributed data. When data did not conform to a normal distribution, a Kruskal-Wallis test was performed, followed by Dunn’s multiple comparisons without any correction. Results are presented as mean ± standard error of mean (SEM) with the n value for each dataset. Statistically significant effects and n numbers are detailed in the corresponding figure legend or STAR Methods. The significance threshold was set at 0.05, two-tailed (not significant, ns, p > 0.05; *p < 0.05; **p < 0.01; ***p < 0.001). No data exclusion was performed. Data for RNA-seq were processed according to the description in the STAR Methods sections, and statistical details are explained in the results and corresponding figure legends.

## Acknowledgements

We thank Belén Andrés and María Aurelia Torregrosa for their technical support. We thank Teresa Guillamón-Vivancos and Miguel Valdeolmillos for their comments and input on the manuscript and other members of López-Bendito’s laboratory for stimulating discussions. This work was supported by grants from the Swiss National Science Foundation (31003A_175776 and 310030_219370) and the Novartis Research Foundation to F.M.R., and by grants from the European Research Council (ERC) under the European Union’s Horizon 2020 research and innovation programme (ERC-2021- ADG-101054313 SPONTSENSE), PID2021-127112NB-I00 from the MCIN/AEI /10.13039/501100011033/ and ERDF A way to make Europe, and Generalitat Valenciana, Conselleria d’Educació, Universitats, i Ocupació (PROMETEO 2021/052) to G.L.-B.

## Author contributions

Conceptualization, M.A-M. and G.L-B.; methodology, M.A-M., G.P., MP.M. and LM.R-M; data curation, M.A-M., G.P., MP.M. and L.P.; transcriptomic analysis, L.P.; writing –original draft, M.A-M., and G.L-B.; writing – review & editing, M.A-M., L.P., G.P., FJ.M., FM.R., and G.L-B.; funding acquisition, FM.R., and G.L-B.; resources, FM.R., and G.L- B.; supervision, FJ.M., FM.R., and G.L-B.

## Competing interests

The authors declare no competing interests.

**Declaration of generative AI and AI-assisted technologies in the writing process** During the preparation of this work the authors used Chat-GPT in order to streamline some parts of the text. After using this tool, the authors reviewed and edited the content as needed and take full responsibility for the content of the publication.

## Resource Availability Lead contact

Additional information and requests for resources and reagents should be directed to the lead contact, Guillermina López-Bendito.

## Materials availability

This study did not generate new mouse lines. This study did not generate new unique reagents.

## Data availability

RNAseq datasets have been deposited at the National Center for Biotechnology Information (NCBI) Gene Expression Omnibus (GEO) and will be publicly available as of the date of publication.

Any additional information required to reanalyze the data reported in this work paper is available from the lead contact upon request.

## Code availability

This study did not generate original code. For *ex vivo* experiments analysis, calcium imaging code previously reported^21^ has been deposited at https://github.com/ackman678/CalciumDX. For *in vivo* calcium imaging experiments, the code from WholeBrainDX repository^60^ (available at https://github.com/ackman678/wholeBrainDX) was employed to prepared the supplementary videos.

**Table S1.** Differential expression analysis of PMBSF vs ALBSF control samples from bulk RNA-seq. Related to Figures 4 and S8.

**Table S2.** Differential expression analysis of EmbWPC vs control-ALBSF samples from bulk RNA-seq. Related to Figures 4 and S8.

**Table S3.** Enrichment analysis of EmbWPC and control-ALBSF samples from bulk RNA-seq. Related to Figures 4 and S9.

## Multimedia files

**Video S1. iDISCO 3D reconstruction of E18 control mouse.** Volume of the whisker pad and upper lip areas in magenta and cyan, respectively. Volume of nice whisker pad and nine upper lip follicles in yellow. Related to Figures 1 and S1.

**Video S2. iDISCO 3D reconstruction of E18 embWPC mouse.** Volume of the whisker pad and upper lip areas in magenta and cyan, respectively. Volume of nice whisker pad and nine upper lip follicles in yellow. Related to Figures 1 and S1.

**Video S3. *In vivo* cortical evoked responses in the control side (ipsilateral to cauterization) of embWPC mouse at E18.** Stimulations of the whisker pad (magenta) and upper lip (cyan). Dashed lines delineate cortical hemisphere, PMBSF and ALBSF putative cortical territories. Speed x3.3. Related to Figure 2.

Video S4. *In vivo* cortical evoked responses in the whisker pad cauterized (WPC) side (contralateral to cauterization) of embWPC mouse at E18. Stimulations of the whisker pad (magenta) and upper lip (cyan). Dashed lines delineate cortical hemisphere, PMBSF and ALBSF putative cortical territories translated from control side stimulations (“PMBSF” and “ALBSF”). Speed x3.3. Related to Figure 2.

**Video S5. *In vivo* cortical evoked responses in the control side (ipsilateral to cauterization) of embWPC mouse at P4.** Stimulations of the whisker pad (magenta) and upper lip (cyan). Dashed lines delineate cortical hemisphere, PMBSF and ALBSF cortical territories. Speed x3.3. Related to Figure 2.

Video S6. *In vivo* cortical evoked responses in the whisker pad cauterized (WPC) side (contralateral to cauterization) of embWPC mouse at P4. Stimulations of the whisker pad (magenta) and upper lip (cyan). Dashed lines delineate cortical hemisphere, PMBSF and ALBSF cortical territories translated from control side stimulations (“PMBSF” and “ALBSF”). Speed x3.3. Related to Figure 2.

Video S7. Spontaneous thalamic activity in the prospective VPM regions (wpVPM and ulVPM) in an acute slice of control mice at E16. **Dashed lines delineate dLG** (dorsolateral geniculate) nucleus, prospective wpVPM and ulVPM regions. Speed x15. Related to Figure 3.

**Video S8. Spontaneous thalamic activity in the prospective VPM regions (wpVPM” and ulVPM) in an acute slice of embWPC mice at E16.** Dashed lines delineate dLG (dorsolateral geniculate) nucleus, prospective wpVPM and ulVPM regions. Speed x15. Related to Figure 3.

**Video S9. Topographic representations in the cortex of *in vivo* evoked responses in control mouse at P4.** Stimulations of the whisker pad (magenta, blue, green, yellow and orange) and upper lip (cyan and white). Dashed lines delineate cortical hemisphere, PMBSF and ALBSF evoked cortical territories. Speed x3.3. Related to Figure 6.

**Video S10. Topographic representations in the cortex of *in vivo* evoked responses in embWPC mouse at P4.** Stimulations of the upper lip (cyan and white). Dashed lines delineate cortical hemisphere and ALBSF evoked cortical territory. Speed x3.3. Related to Figure 6.

**Figure S1.**
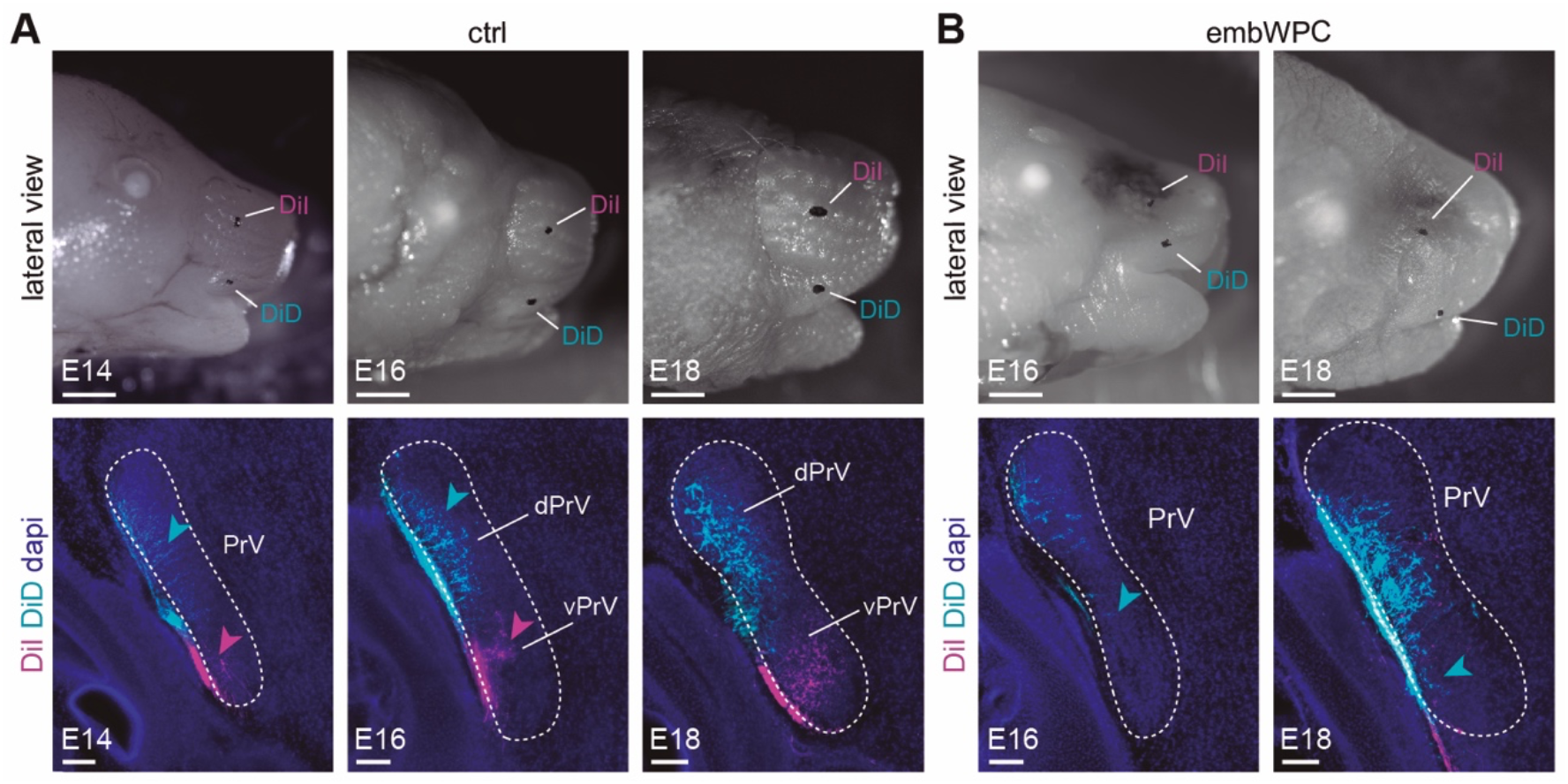
Early reorganization of peripheral axons at the trigeminal nucleus, Related to. Figure 1**. (A)** Upper panels, lateral views of control mouse heads showing DiI and DiD crystal placements in the whisker pad and upper lip, respectively, at distinct developmental stages (E14, E16 and E18). Lower panels, coronal sections showing labelled axons at the vPrV (magenta, arrowheads) and dPrV (cyan, arrowheads) (n= 6 mice at E14, n= 9 mice at E16, n= 6 mice at E18). **(B)** Upper panels, lateral views of embWPC mouse heads showing DiI and DiD crystal placements in the whisker pad and upper lip, respectively, at distinct developmental stages (E16 and E18). Lower panels, coronal sections showing displaced upper lip axons at the vPrV (cyan, arrowheads) (n= 9 embWPC mice at E16, n= 5 embWPC mice at E18). E, embryonic; embWPC, embryonic whisker pad cauterized; PrV, principal nucleus; vPrV, ventral principal nucleus; dPrV, dorsal principal nucleus; DiI, 1,1′-dioctadecyl 3,3,3′,3′- tetramethylindocarbocyanine perchlorate; DiD, 1,1’-Dioctadecyl-3,3,3’,3’- Tetramethylindodicarbocyanine, 4-Chlorobenzenesulfonate. Scale bars, (A, top) and (B) 1000 μm; (A, bottom) and (B) 100 μm.

**Figure S2.**
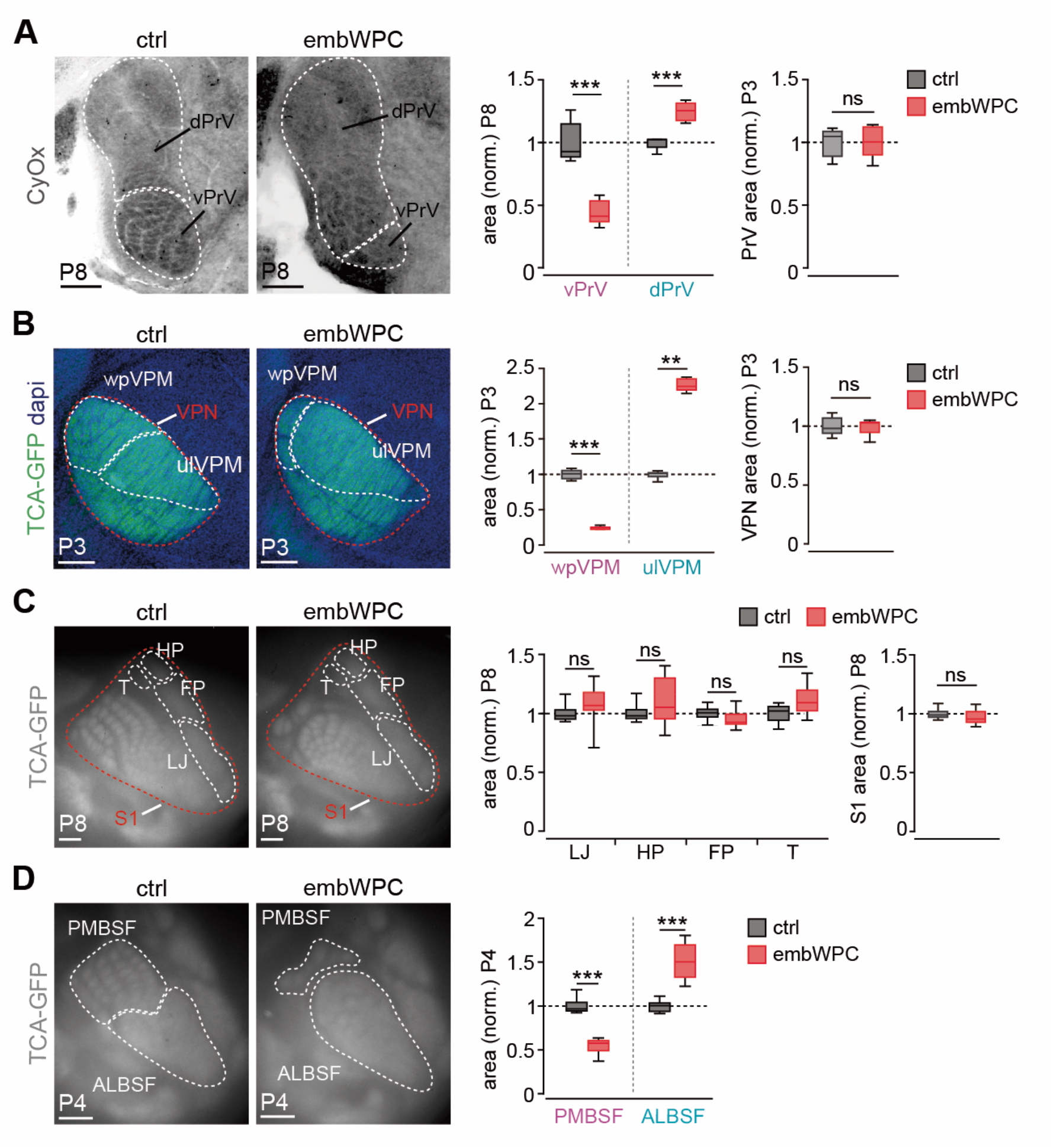
Intra-modal rescaling of facial whiskers representations in cortical and subcortical territories in embWPC mice. Related to. Figure 1**. (A)** Coronal sections of the trigeminal PrV nucleus stained with CyOx in control and embWPC at P8. Quantification of the data (vPrV and dPrV areas, n= 5 ctrl mice, n= 5 embWPC mice, unpaired two-tailed Student *t*-test; PrV area, n= 5 ctrl mice, n= 5 embWPC mice, unpaired two-tailed Student *t*-test. ns, p > 0.9). **(B)** Coronal sections showing TCA-GFP labelling in the PMBSF, ALBSF and VPN of the thalamus in control and embWPC at P3. Quantification of the data (n= 5 ctrl mice, n= 5 embWPC mice, PMBSF, unpaired two- tailed Student *t*-test. ALBSF, Mann-Whitney *U*-test; VPN at P3, n= 5 ctrl mice, n= 5 embWPC mice, Mann-Whitney *U*-test. ns, p > 0.99). **(C)** Surface view of thalamocortical terminals (TCA-GFP+) in the PMBSF and ALBSF in control and embWPC mice at P8. Quantification of the data (n= 8 ctrl mice, n= 8 embWPC mice, unpaired two-tailed Student *t*-test. ns, p > 0.05. S1 area, n= 8 ctrl mice, n= 9 embWPC mice, unpaired two- tailed Student *t*-test. ns, p= 0.29). **(D)** Surface view of thalamocortical terminals (TCA-GFP+) in the PMBSF and ALBSF in control and embWPC mice at P4. Quantification of the data (n= 6 ctrl mice, n= 6 embWPC mice, unpaired two-tailed Student *t*-test; ***p < 0.001). CyOx, Cytochrome oxidase; P, postnatal; wpVPM, whisker pad recipient ventral posteromedial nucleus; ulVPM, upper lip recipient ventral posteromedial nucleus; VPN, ventroposterior nucleus; TCA-GFP, thalamocortical axons labelled with green fluorescent protein; S1, primary somatosensory cortex; HP, hindpaw; FP, forepaw; LJ, lowerjaw; T, trunk; PMBSF, postero-medial barrel subfield; ALBSF, antero-lateral barrel subfield; norm., normalized. Scale bars, (A) and (B) 200 μm; (C) and (D) 500 μm. Boxplots show the medians with the interquartile range (box) and range (whiskers). ns, not significant. **p < 0.01, ***p < 0.001.

**Figure S3.**
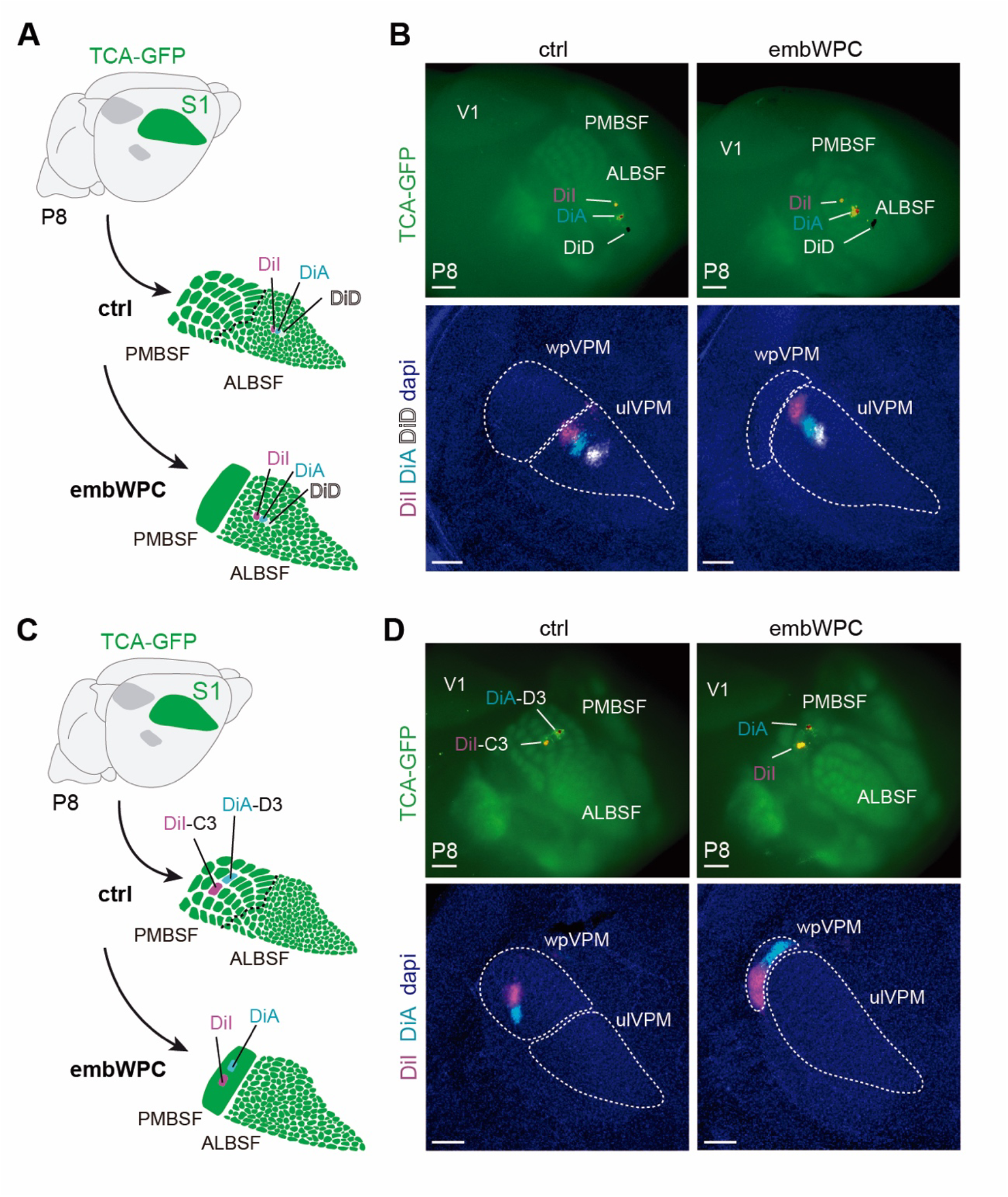
Thalamocortical point-to-point topography in embWPC mice. Related to. Figure 1**. (A)** Schema representing the experimental paradigm. **(B)** Upper panels, DiI, DiA and DiD crystal placements subsequent non-overlapping points in the ALBSF in control and embWPC at P8. Lower panels, backlabelled cells at the ulVPM showed a shifted relative displacement toward the wpVPM in embWPC mice (n= 8 ctrl mice, n= 8 embWPC mice). **(C)** Schema representing the experimental paradigm. **(D)** Upper panels, DiI and DiA crystal placements in the C3 and D3 cortical barrels, respectively, in PMBSF control and in the remaining territory of the PMBSF in embWPC at P8. Lower panels, backlabelled cells maintain their point-to-point distribution at the wpVPM both in control and embWPC mice (n= 9 ctrl mice, n= 9 embWPC mice). V1, primary visual cortex; DiA, 4-[4-(dihexadecylamino) styryl]-N-methylpyridinium iodide. Scale bars, (B and D, top) 500 μm; (B and D, bottom) 200 μm.

**Figure S4.**
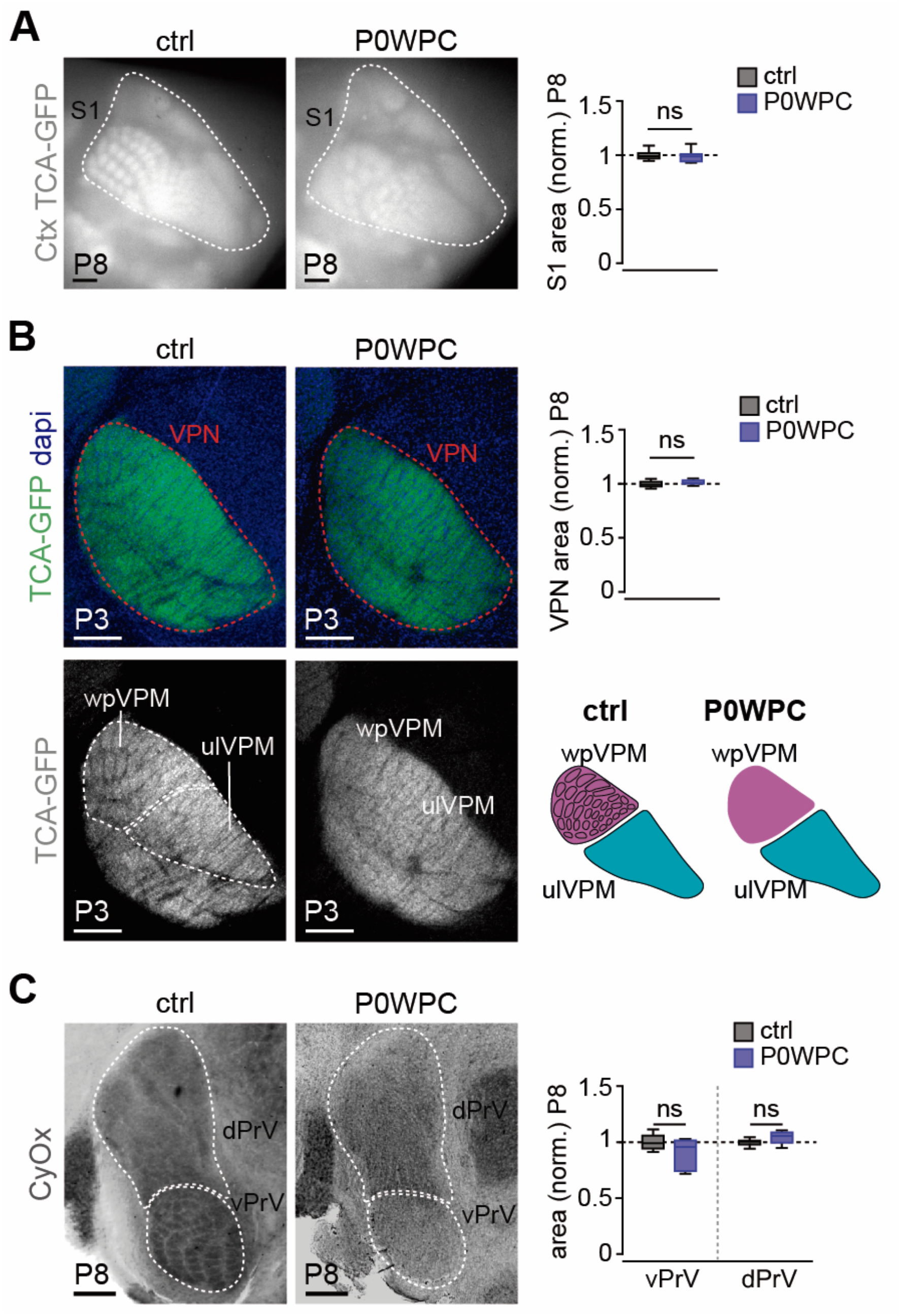
Cortical and subcortical areas for facial whiskers representations remained unchanged in P0WPC mice. Related to. Figure 1**. (A)** Surface view of thalamocortical terminals (TCA-GFP+) in the cortical S1 in control and P0WPC mice at P8. Quantification of the total S1 area (n= 8 ctrl mice, n= 9 P0WPC mice, unpaired two-tailed Student *t*-test. ns, p= 0.66). **(B)** Coronal sections showing TCA-GFP labelling in the PMBSF, ALBSF and VPN of the thalamus in control and P0WPC at P3. Quantification of the VPN area and schema representing the results (n= 8 ctrl mice, n= 9 P0WPC mice, unpaired two-tailed Student *t*-test. ns, p= 0.39). **(C)** Coronal sections of the trigeminal PrV nucleus stained with CyOx in control and P0WPC at P8. Quantification of the vPrV and dPrV area (n= 5 ctrl mice, n= 5 P0WPC mice, unpaired two-tailed Student *t*-test. ns, p> 0.20). P0WPC, postnatal day 0 whisker pad cauterized. Scale bars, (A) 500 μm; (B) and (C) 200 μm. Boxplots show the medians with the interquartile range (box) and range (whiskers). ns, not significant.

**Figure S5.**
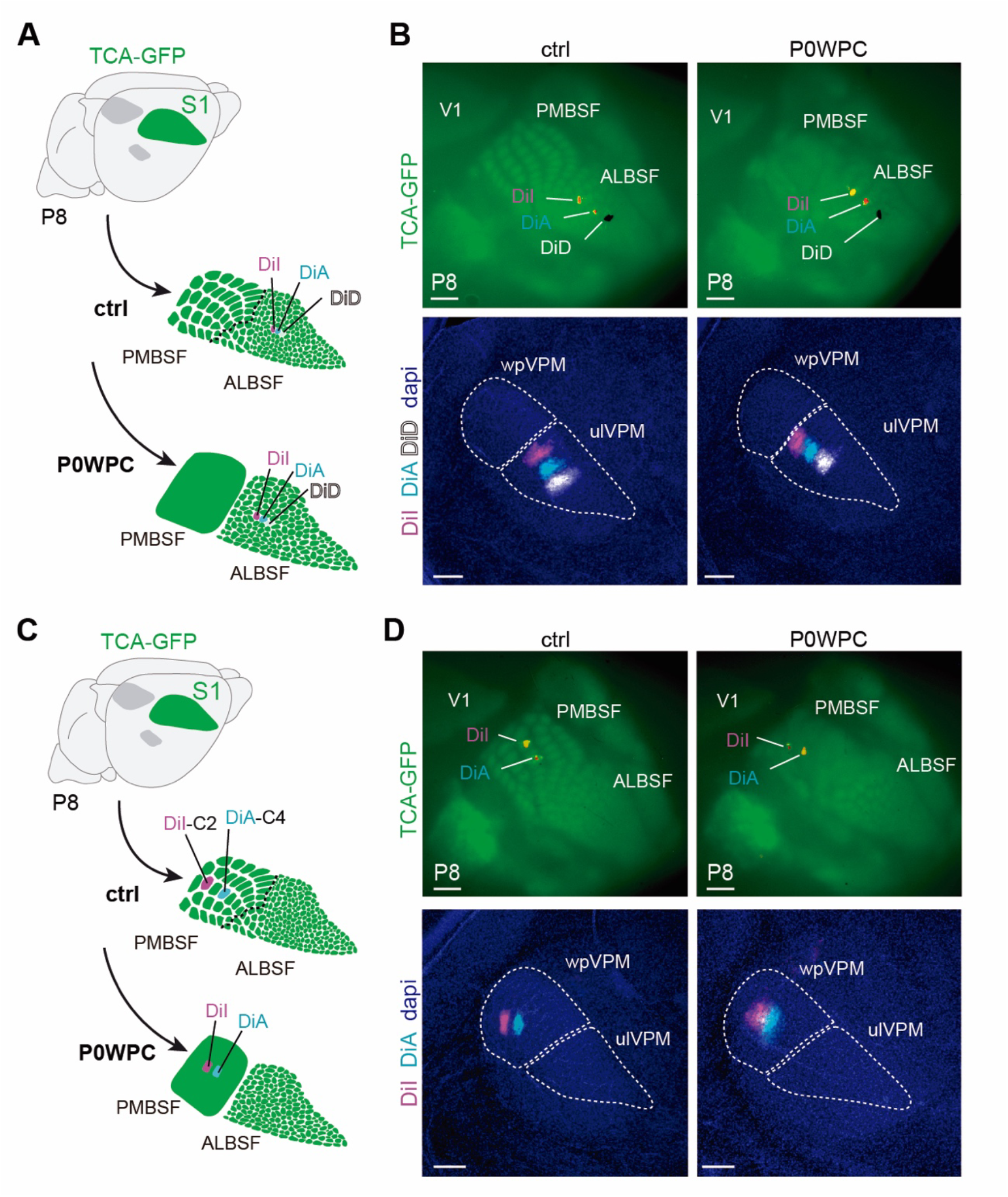
Thalamocortical point-to-point topography in P0WPC mice. Related to. Figure 1**. (A)** Schema representing the experimental paradigm. **(B)** Upper panels, DiI, DiA and DiD crystal placements subsequent non-overlapping positions in the ALBSF in control and P0WPC at P8. Lower panels, backlabelled cells at the ulVPM (n= 7 ctrl mice, n= 7 P0WPC mice). **(C)** Schema representing the experimental paradigm. **(D)** Upper panels, DiI and DiA crystal placements in the C2 and C4 cortical barrels, respectively, in control and in the deprived PMBSF territory in P0WPC at P8. Lower panels, backlabelled cells showing their point-to-point distribution at the wpVPM in control and P0WPC mice (n= 7 ctrl mice, n= 7 P0WPC mice). Scale bars, (B and D, top) 500 μm; (B and D, bottom) 200 μm.

**Figure S6.**
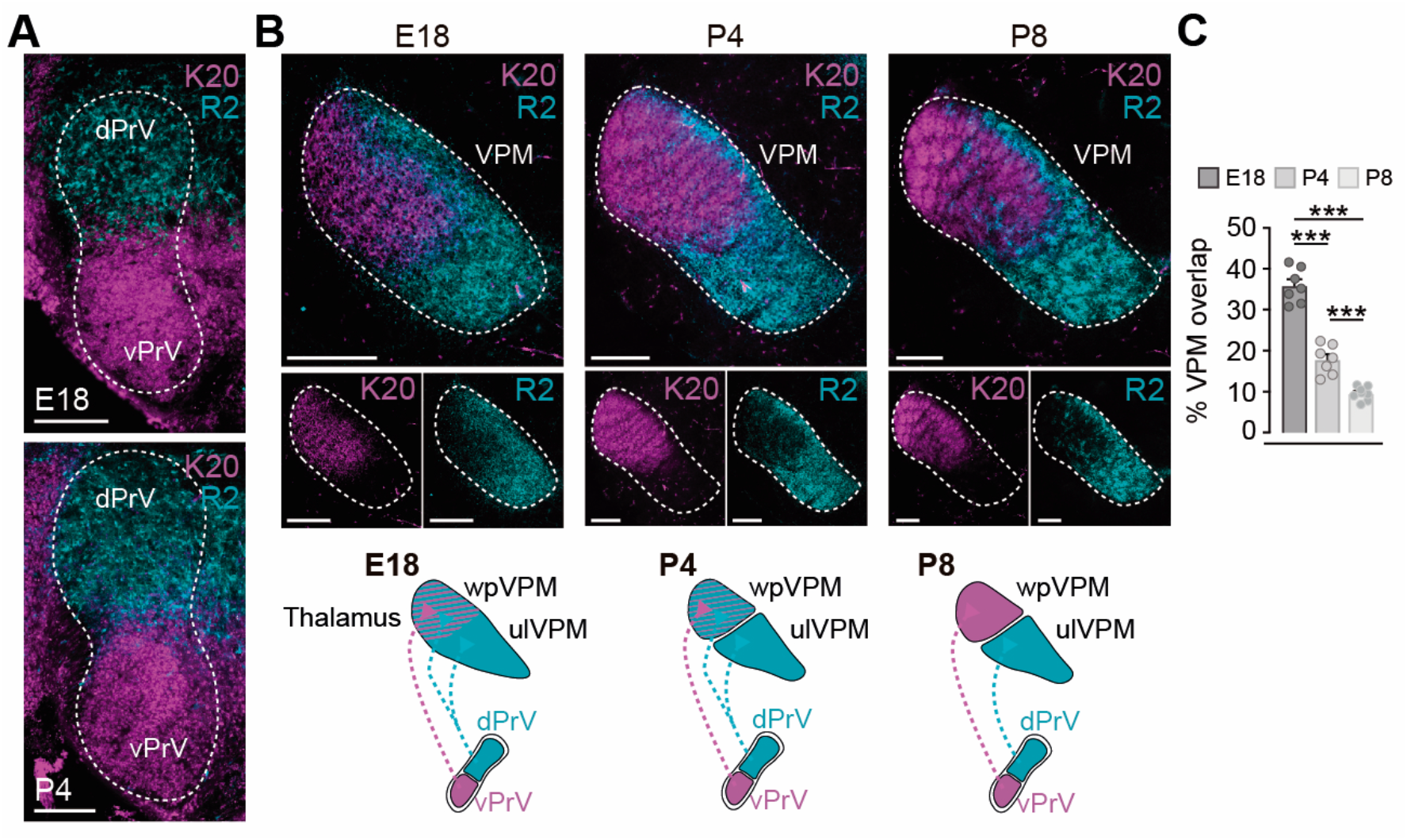
Developmental progression of trigemino-thalamic axons. Related to. Figure 3**. (A)** Coronal sections from the *Krox20-zsgreen::R2-mCherry* double transgenic mouse showing cells expressing Krox20 (magenta) at the vPrV and R2 (cyan) at the dPrV at E18 and P4 (n= 7 E18, n= 7 P4). **(B)** Coronal sections from the *Krox20- zsgreen::R2-mCherry* double transgenic mice showing Krox20-labelled axons (magenta) and R2-labelled axons (cyan) at the VPM of the thalamus at E18, P4 and P8. Schemas representing the results found. **(C)** Quantification of the percentage of developmental overlap between vPrV and dPrV axons at the thalamus (n= 7 E18, n= 7 P4, n= 8 P8, Two-way ANOVA test: ***p <0.001. Tukey’s multiple comparison test *post-hoc* analysis). Scale bars, 200 μm. Bar graphs show the means ± SEM. ***p <0.001.

**Figure S7.**
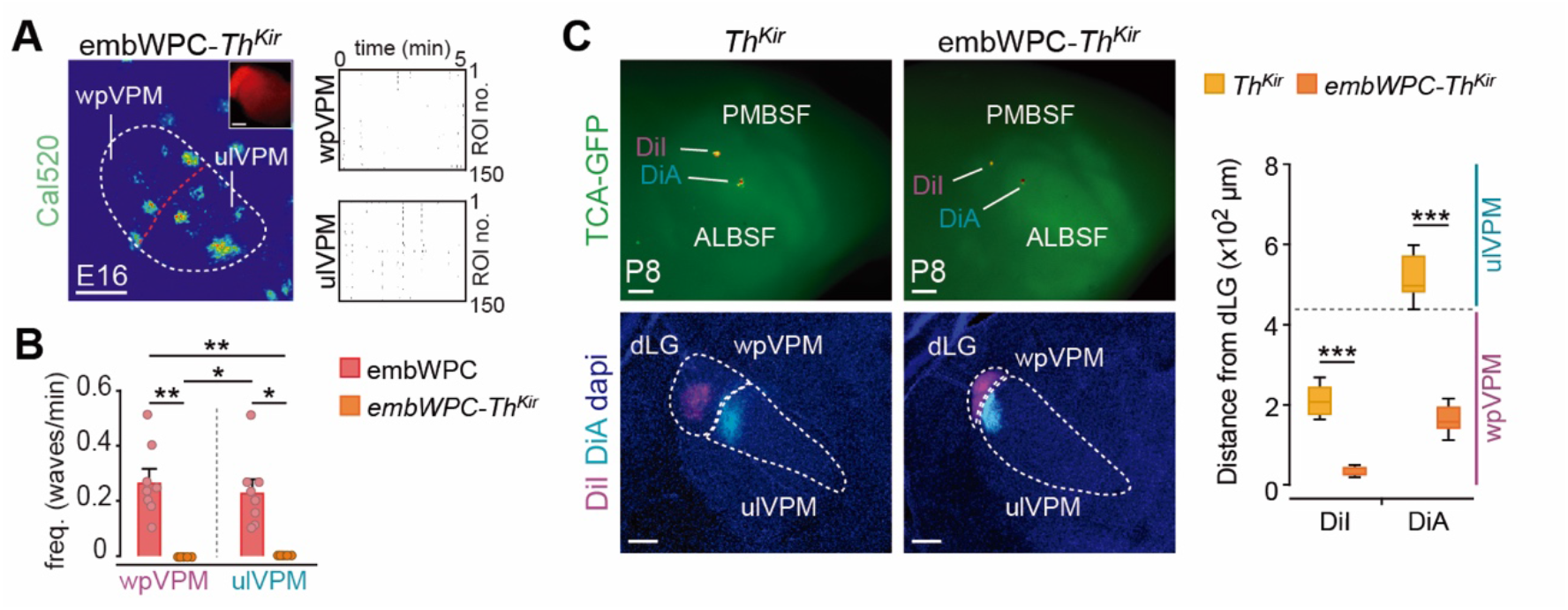
Thalamocortical point-to-point topography is shifted in embWPC-*Th^Kir^* but not in *Th^Kir^* mouse. Related to. Figure 3**. (A)** Maximal projection of *ex vivo* spontaneous calcium activity in the prospective VPM sub-regions (wpVPM, wp input- receiving neurons, and ulVPM, ul input-receiving neurons) of the thalamus in acute slices at E16. Raster plots of 5 minutes. **(B)** Quantification of the frequency of waves activity in the VPM in both embWPC and embWPC-*Th^Kir^* mice (n= 8 embWPC mice, n= 5 embWPC-*Th^Kir^* mice, Kruskal-Wallis test, ***p <0.001. Dunn’s multiple comparison test post-hoc analysis). **(C)** Upper panels, DiI and DiA crystal placements in the cortical PMBSF and ALBSF areas, respectively. Lower panels, backlabelled cells at the thalamus show shifted positions within the ALBSF thalamic area in the embWPC-*Th^Kir^* mice as compared to *Th^Kir^* at P8. Quantification of the position of backlabelled cells with respect to the distance to the dLG nucleus. The gray horizontal dashed line in the graph represents the separation between PMBSF and ALBSF (n= 6 *Th^Kir^* mice, n= 6 embWPC- *Th^Kir^* mice, unpaired two-tailed Student *t*-test). Scale bars, (A) and (C, bottom) 200 μm; (C, top) 500 μm. Bar graphs show the means ± SEM. *p < 0.05, **p < 0.01, ***p < 0.001.

**Figure S8.**
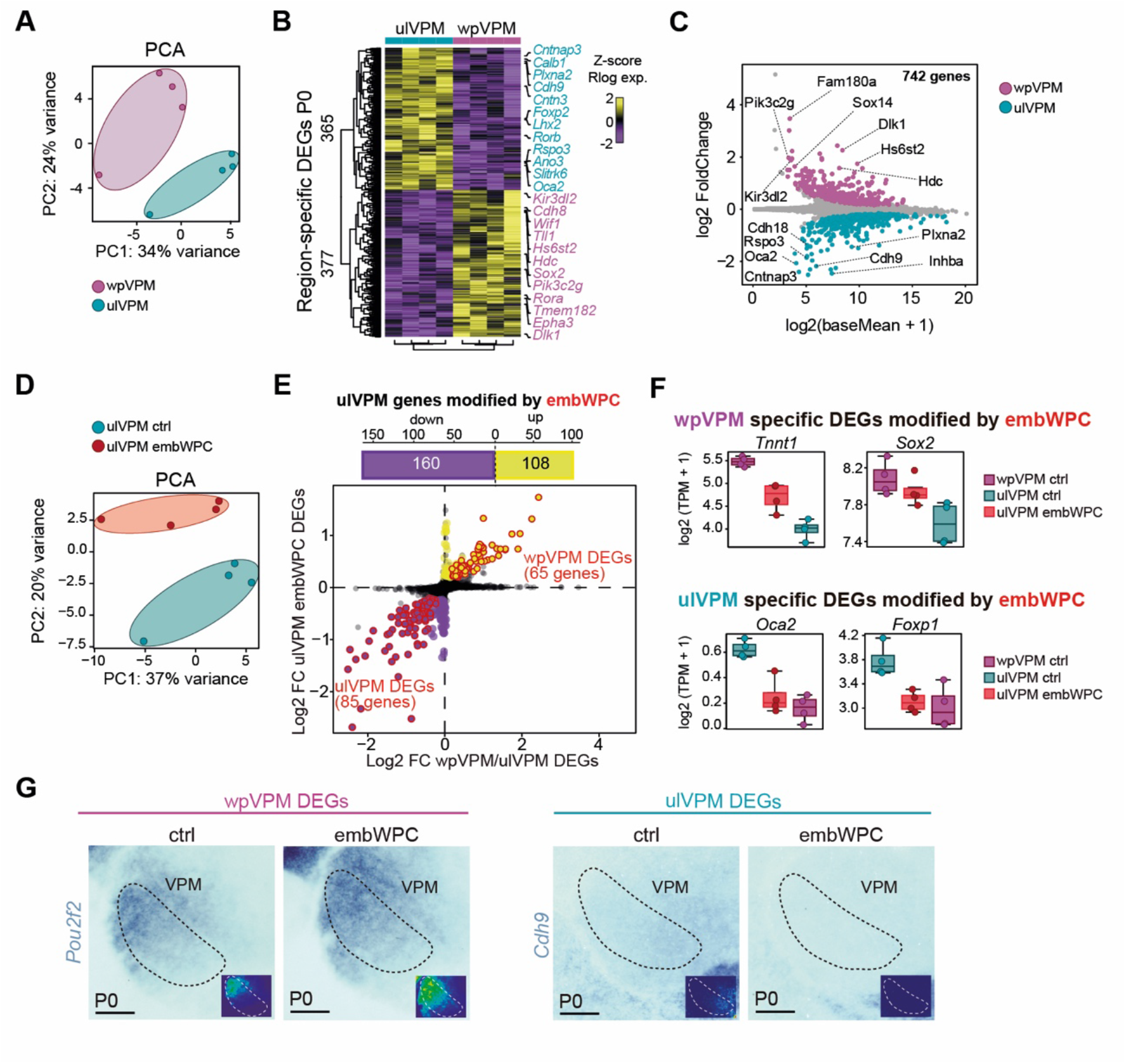
Region-specific transcriptional signatures in the thalamic VPM sub- regions in control and embWPC mice at P0. Related to. Figure 4**. (A)** Principal Component Analysis (PCA) of wpVPM (n= 4) and ulVPM (n= 4) control mice at P0. **(B)** Heatmap of normalized regularized logarithm (Rlog) Z-score of expression and unbiased clustering of ulVPM- and wpVPM-specific Differentially Expressed Genes (DEGs) at P0. The color-code (yellow, high expression; purple, low expression) corresponds to the log2FC. **(C)** MA plot showing the log2 FoldChange and the mean expression distribution of DEGs. The magenta and cyan dots represent wpVPM and ulVPM DEGs, respectively, with the top 7 protein coding genes listed in every region. **(D)** Principal Component Analysis (PCA) of thalamic control ulVPM (n= 4) and embWPC ulVPM (n= 4) mice at P0. **(E)** Top, histogram showing ulVPM DEGs that are up- and downregulated by embWPC at P0. Bottom, scatterplot of log2Foldchange between ulVPM DEGs modified by embWPC and wpVPM versus ulVPM region-specific DEGs. **(F)** Boxplots displaying TPM expression levels of selected wpVPM DEGs and ulVPM DEGs modified in the ulVPM of the embWPC mice. Boxplots show the medians with the interquartile range (box) and range (whiskers). **(G)** Coronal sections showing by in situ hybridization the change in the pattern of expression of additional wpVPM and ulVPM DEGs in the embWPC mouse at P0 (n= 5 ctrl, n= 5 embWPC, for each probe). Scale bars, 200 μm.

**Figure S9.**
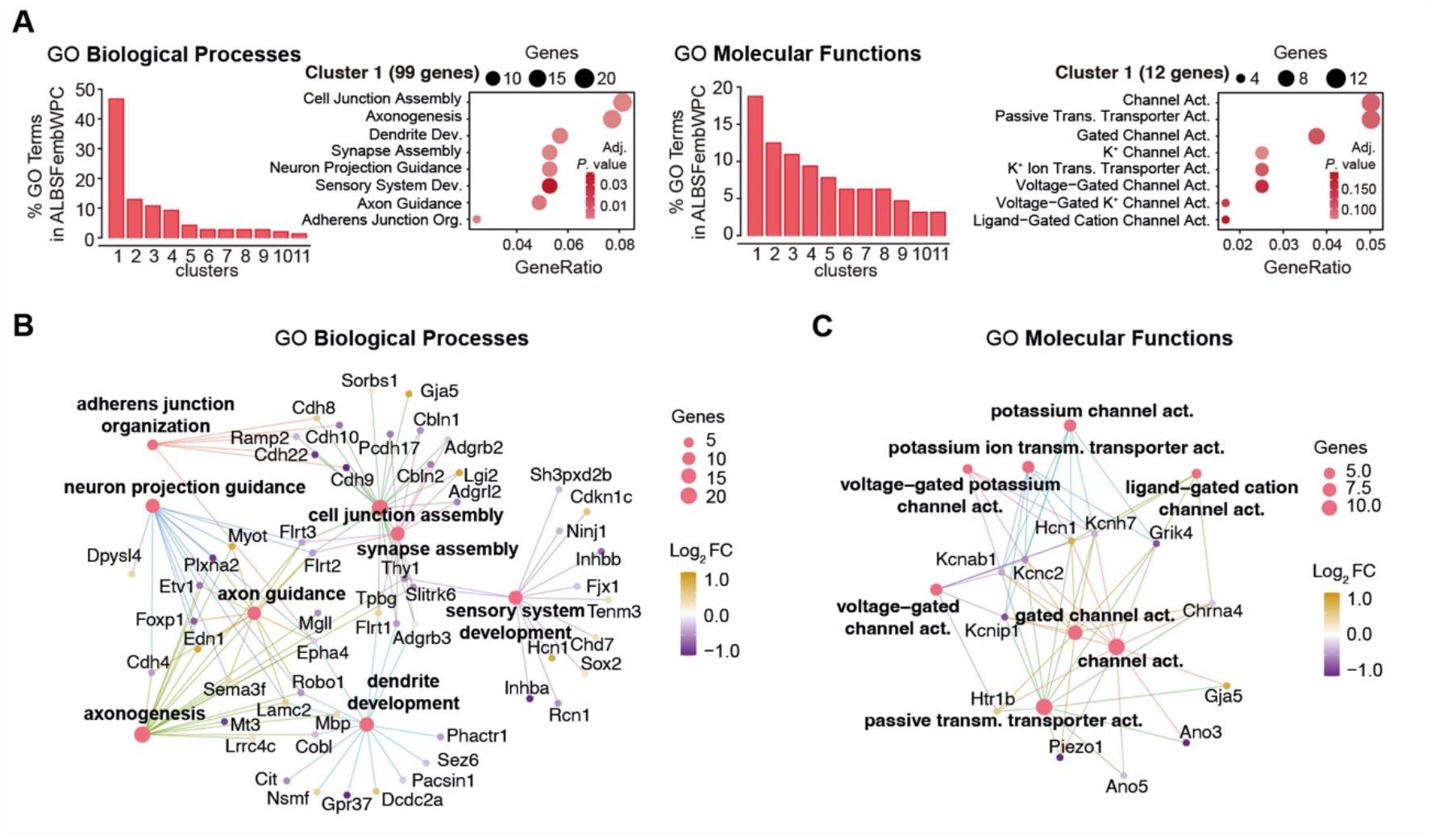
Gene Ontology analysis of region-specific genes modified in embWPC mice. Related to. Figure 4**. (A)** Gene Ontology (GO) functional analysis (Biological Processes, left; and Molecular Functions, right) showing the top ranked clusters of GO terms enriched in the region-specific gene-set modified in embWPC-ulVPM. **(B)** Gene ontology (GO) biological process (BP) enrichment analysis of wpVPM and ulVPM DEGs modified by embWPC in ulVPM. **(C)** Gene ontology (GO) molecular function (MF) enrichment analysis of wpVPM and ulVPM DEGs modified by embWPC in ulVPM. The size of every node (enriched term) in the gene networks represents the number of genes enriched and the color-code (yellow, high expression; purple, low expression) corresponds to the log2FC in the wpVPM vs ulVPM Differential expression analysis (DEA).

